# Identification of novel and potent inhibitors of SARS-CoV-2 main protease from DNA-encoded chemical libraries

**DOI:** 10.1101/2024.03.16.585341

**Authors:** Dario Akaberi, Monireh Pourghasemi Lati, Janina Krambrich, Julia Berger, Grace Neilsen, Emilia Strandback, S. Pauliina Turunen, Johan Wannberg, Hjalmar Gullberg, Martin Moche, Praveen Kumar Chinthakindi, Tomas Nyman, Stefan G. Sarafianos, Anja Sandström, Josef D. Järhult, Kristian Sandberg, Åke Lundkvist, Oscar Verho, Johan Lennerstrand

## Abstract

In vitro screening of large compounds libraries with automated high-throughput screening is expensive, time consuming and requires dedicated infrastructures. Conversely, the selection of DNA-encoded chemical libraries (DECL) can be rapidly performed with routine equipment available in most laboratories. In this study we identified novel inhibitors of SARS-CoV-2 main protease (M^pro^) through the affinity-based selection of the DELopen library (open access for academics), containing 4.2 billion compounds. The identified inhibitors were peptide-like compounds containing an N-terminal electrophilic group able to form a covalent bond with the nucleophilic Cys145 of M^pro^, as confirmed by x-ray crystallography. This DECL selection campaign enabled the discovery of the unoptimized compound SLL11 displaying an IC_50_ of 30 nM, proving that the rapid exploration of large chemical spaces enabled by DECL technology, allows for the direct identification of potent inhibitors avoiding several rounds of iterative medicinal chemistry. Compound MP1, a close analogue of SLL11, showed antiviral activity against SARS-CoV-2 in the low micromolar range when tested in Caco-2 and Calu-3 (EC_50_ = 2.3 *µ*M) cell lines. As peptide-like compounds can suffer from low cell permeability and metabolic stability, the cyclization of the compounds as well as the substitution of selected residues with D-enantiomers will be explored in the future to improve the antiviral activity of these novel compounds.

## 1 Introduction

Four years after the COVID-19 pandemic started, infections are driven by the emergence of new SARS-CoV-2 variants of concern (1). Although mRNA vaccines have been instrumental in reducing severe disease and hospitalization, achieving long-term immunity appears challenging and periodical boosting is required. The now dominant Omicron subvariants present over 30 mutations in the spike protein (S gene), conferring resistance to neutralizing antibodies induced by previous mRNA vaccines, bivalent vaccines or infection with a previous variant (2–4). Also the efficacy of monoclonal antibodies used to prevent SARS-CoV-2 infection (5), disease progression and death (6, 7) can be reduced by emerging variants presenting new mutations in the spike gene (6, 7). In this scenario, antiviral options for the prevention and treatment of SARS-CoV-2 infection in immunosuppressed and high-risk subjects are necessary to further reduce the COVID-19 burden. Moreover, broad-spectrum antivirals active against several coronavirus will be instrumental for preventing or mitigating the next pandemic by reducing early transmission and providing a starting point for the development of more potent compounds if necessary.

SARS-CoV-2 3-chymotrypsin-like cysteine protease (3CL protease) also known as main protease (M^pro^) (8) mediates the maturation of viral proteins by cleaving the two viral polyproteins pp1a and pp1ab at 11 sites (9). M^pro^ represents an attractive target for the development of antivirals against SARS-CoV-2 as it is essential for the viral life cycle and structurally conserved among alpha and beta coronaviruses (10), allowing the development of potent pan-coronavirus protease inhibitors (11–13). The feasibility of M^pro^ inhibitors as prophylaxis or treatment against SARS-CoV-2 infection has also already been proven. Two orally administered M^pro^ inhibitors, nirmatrelvir (approved by the FDA and EMA) and ensitrelvir (approved in Japan by the Ministry of Health, Labour and Welfare (MHLW) (14) are currently approved for emergency treatment of COVID-19.

SARS-CoV-2 inhibitors have been identified through the development of substrate derived peptide-like compounds (8, 15, 16) as well as screening of large libraries of compounds and fragments using different techniques such as *in silico* screening (11), high-throughput screening (17) and crystallographic screening (18).

Two studies have also reported the identification of novel M^pro^ inhibitors, with IC_50_ and EC_50_ in the low micromolar range (19, 20), using DNA encoded chemical libraries (DECLs). DECLs are large collections of compounds tagged with a unique DNA barcode that are screened by affinity to select binders for a target of interest (21). Following the selection process, the chemical structures of the binders are elucidated through the sequencing of their unique DNA tags. This technology is particularly attractive because the combinatorial nature of DECL combined with affinity screening allow for rapid and cost-efficient exploration of large portions of chemical space *in vitro*.

Here we report a new class of peptide-like inhibitors of M^pro^ identified using the commercially available DELopen platform (Wuxi Apptech) that provides a library of 4.2 billion compounds (screened by the user) as well as the services necessary for the sequence to structure decoding of binders’ DNA tags. The most potent compound (MP1) inhibited the activity of SARS-CoV-2 M^pro^ with IC_50_ = 24 nM and SARS-CoV-2 infection with EC_50_ = 2.3 *µ*M in cell-based assays.

## 2 Material and methods

### 2.1 Bead capture test

Immobilization of M^pro^-Avi to Dynabeads MyOne Streptavidin T1 paramagnetic beads was tested before affinity selection according to a protocol provided with the DELopen kit. All reagents were added to 1.5 ml Epperdorf DNA Low Binding tubes during bead capture test and affinity selection. In brief, paramagnetic beads were washed three times with 1X wash buffer (50 mM Tris-HCl pH 7.5 – 150 mM sodium chloride – 0.05% Tween-20 – 1 mM dithiothreitol) using DynaMag-2 magnetic stand (Invitrogen). M^pro^-Avi (6 *µ*g) was diluted to 1X selection buffer (50 mM Tris-HCl pH 7.5 – 150 mM sodium chloride – 0.05% Tween-20 – 1 mM dithiothreitol - 0.1 mg/ml sheared salmon sperm DNA) and an aliquot was collected as input sample. Remaining Avi-tagged protein (5 *µ*g) was immobilized to Dynabeads MyOne Streptavidin T1 paramagnetic beads at room temperature for 30 minutes. Flowthrough was collected and beads were washed once with selection buffer. Beads were suspended to 1X selection buffer, a ‘beads’ sample was collected, and the remaining bead suspension was heated at 95° C for 10 minutes. Beads were collected using a magnetic stand and solution was collected as heated eluate sample. Beads were resuspended to selection buffer (heated beads). To visualize the relative amount of immobilized and eluted protein the bead capture test samples were separated by SDS-PAGE. Samples were denatured and reduced using NuPAGE LDS Sample Buffer (4X) (Invitrogen) and NuPAGE Sample Reducing Agent (10X) (Invitrogen) by heating at 95° C for 5 minutes. Samples were separated on NuPAGE 4-12% Bis-Tris Mini Protein Gels (Invitrogen) using matched electrophoresis run chamber at 200V for 35 minutes. Proteins on the gels were visualized using InstantBlue Coomassie Protein Stain (Abcam), destaining with distilled water.

### 2.2 DECL affinity selection

Three rounds of affinity selection were performed using the 3rd generation DELopen DECL (WuXi Apptec; hits.wuxiapptec.com/delopen) according to the provider’s instructions. Before each round, M^pro^-Avi (5 *µ*g) was immobilized to Dynabeads MyOne Streptavidin T1 paramagnetic beads at room temperature for 30 minutes in 1X selection buffer. Reversible small molecule inhibitors were added at 20 *µ*M concentration for the last 10 minutes of immobilization and included into DEL incubation mixtures at 20 *µ*M final concentration. Covalent inhibitors at 5 *µ*M were preincubated with immobilized target for 10 minutes at room temperature but not added to the DEL incubation. Bead-immobilized target at final concentration 1.7 *µ*M was incubated with the DELopen compound library in 1X selection buffer (50 mM Tris-HCl pH 7.5 – 150 mM sodium chloride – 0.05% Tween-20 – 1 mM dithiothreitol - 0.1 mg/ml sheared salmon sperm DNA) for one hour at room temperature with gentle rotation. Three wash cycles were performed with 1X selection buffer (50 mM Tris-HCl pH 7.5 – 150 mM sodium chloride – 0.05% Tween-20 – 1 mM dithiothreitol) after each round. The compound release method was heating for 10 minutes at 98°C after each round. Samples collected after the third selection round were stored at -80°C before transfer to Wuxi App Tec for post-selection quality control and deep-sequencing.

### 2.3 Protease expression and purification

SARS-CoV-2 3CL protease (M^pro^) used for enzymatic assays and co-crystallization experiments was produced as previously described (22). The used construct contained nucleotide sequences corresponding to SARS-CoV-2 M^pro^ residues S1-Q306 (Chinese isolate, NCBI accession number YP_009725301). A detailed protocol is provided in the supporting information.

Avi-tagged SARS-CoV-2 M^pro^ (batch nr. MPRO_p009 GP-AVI) protease used for affinity selection of the DECL was kindly provided by Martin Walsh, Diamond Light Source, UK. A detailed protocol for the expression and purification of the enzyme is provided in the supporting information.

SARS-CoV-2 M^pro^ Washington strain (WA1, accession number MT246667), WT (M^pro^-WT) or carrying the E166V mutation (M^pro^-E166V) was expressed and purified as described in (23).

### 2.4 Protein-ligand co-crystallization and x-ray data collections

Compounds SLL11, SLL12 and MP9 were added at 27-fold, 34-fold and 13-fold excess respectively to a 5.2 mg/ml SARS-CoV-2 protease solution in 20 mM HEPES (pH 7.5) and 50 mM NaCl. Before co-crystallization, the non-dissolved ligand was spun down by centrifugation 13,000 rpm for 30 seconds in a Hettich 200 R microcentrifuge.

For ligand SLL11 sitting-drop co-crystallization in 96-well Corning 3550 plates was performed by mixing equal amounts (150 + 150 nl) of ligand incubated protein with a solution consisting of 18% w/v PEG3350, 0.2M potassium-thiocyanate and 0.1M bis-tris propane pH 8.5 using a mosquito robot. The first crystals of the SLL11 complex appeared after 53 days in the well solution and were harvested on day 98. For ligand SLL12 we performed hanging-drop co-crystallization in 24-well VDXm plates by manual mixing equal amounts (1+1 μl) of ligand incubated protein with a solution consisting of 20% w/v PEG3350, 0.2M potassium-thiocyanate and 0.1M bis-tris propane pH 8.5. The first crystals of the M^pro^-SLL12 complex appeared after 35 days and was harvested at day 98. For ligand MP9 we performed sitting-drop co-crystallization in 96-well MRC plates by mixing equal volumes 150 + 150 nl of ligand incubated protein with a solution consisting of 18% w/v PEG3350, 0.2M potassium-thiocyanate and 0.1M bis-tris propane pH 8.7 using a mosquito robot. The first crystals of the MP9 complex appeared after 5 days and were harvested at day 9. All crystals were cryo protected by adding cryo solution consisting of a solution supplemented with 15-30% Glycerol and 50mM NaCl to the crystal droplet right before the crystals were picked up using Dual-Thickness MicroLoop and flash-frozen in liquid nitrogen.

X-ray diffraction datasets were collected at cryogenic 100 K temperature, using wavelength 0.9763 Å, at MAX IV BioMAX beamline (24) in Lund Sweden (Compounds: SLL11, PDB ID 9EO6, and SLL12, PDB ID 9EOR) and, using wavelength 0.9795 Å, at Diamond Light Source (25), i04 beamline in Oxfordshire, UK (Compound MP9, PDB ID 9EOX). All complex datasets crystallize in space group P1 with six SARS-CoV2 M^Pro^ molecules in the asymmetric unit and we collected 360 degrees of data using a single crystal for each complex and processed our datasets using XDS (26) part of XDSAPP (27). The crystals diffracted better in some directions, and we therefore applied elliptical data truncation using the staraniso webserver (staraniso.globalphasing.org) leading to capture of best diffraction data at the expense of completeness in the highest resolution shells.

The structures were solved by molecular replacement using our in-house determined apo structure (7PFL) as search model. For data scaling, molecular replacement and refinement software from the CCP4 suite (28) was used such as aimless (29), phaser (30), refmac5 (31) and coot (32). Refinement dictionaries for the ligands were generated from a ligand SMILES string using the grade web server (grade.globalphasing.org), Acedrg (33) and other recently developed ligand tools (34) in CCP4 to model the ligand-protein covalent bond. We used non-crystallographic symmetry (NCS) restraints throughout refinement that ended when R/Rfree values did not improve further. The structures of compounds SLL11, SLL12 and MP9 have 97.0/96.2/96.5 % residues favored, 2.7/3.3/2.7 % residues allowed, and 0.3/0.5/0.8 % residues in outlier regions of the Ramachandran plot. The SLL11, SLL12 and MP9 structure have been deposited in the protein data bank with accession 9EO6, 9EOR and 9E0X respectively with data collection and refinement statistics presented in Supplementary Table1.

### 2.5 In vitro enzymatic assay

All experiments were carried out as previously reported (22) in black flat-bottomed 96-well plates (Nunc, Thermo Fisher Scientific) in a final volume of 100 *µ*l. Different concentrations of the compounds were incubated with recombinant SARS-CoV-2 protease (final concentration 100nM) in assay buffer (20 mM HEPES pH 7.5, 0.01% Triton X-100) for 10 minutes at room temperature. The enzymatic reaction was started by addition of the substrate DABCYL-Lys-Thr-Ser-Ala-Val-Leu-Gln-Ser-Gly-Phe-Arg-Lys-Met-Glu-EDANS at a final concentration of 20 *µ*M. The fluorescent emission was monitored every 60 seconds for 40 minutes using a Tecan infinite M200 PRO plate reader (Tecan Trading AG, Switzerland) with the excitation wavelength set to 355 nm and the emission wavelength set to 538 nm.

For the initial screening, compounds stocks (10 mM in 100% DMSO) were diluted to 10*µ*M in assay buffer and then further diluted ten times to the final concentration of 1 *µ*M by transferring 10*µ*l in the assay wells (final volume 100 *µ*l). The final concentration of DMSO was kept to 0.01% (v/v) in all wells comprising the controls wells. All compounds and controls were tested with triplicates.

For the SAR study, compound’s IC_50_ were determined with a 12-points concentration series composed of one series of six 1:5 dilutions ranging from 4 to 0.00128 *µ*M (final concentration in the well), and a second series of six 1:5 dilutions ranging from 2 to 0.00064 *µ*M (final concentration in the well). Average IC_50_ and standard error of the means were calculated from two independent experiments with each compound concentration tested in triplicates. DMSO concentrations were always kept ≤ 0.04% (v/v) in all wells.

For resistance testing, MP6, MP9, and nirmatrelvir were tested at final concentrations ranging from 4 *µ*M to 0.00005 *µ*M (eight 1:5 serial dilutions). The recombinant SARS-CoV-2 M^pro^ (WA1, Washington strain) carrying the E166V mutation (M^pro^-E166V) was used at a final concentration of 500nM. The same compounds were also tested at concentration ranging from 100 *µ*M to 0.78 *µ*M (eight 1:2 serial dilutions) against the M^pro^-WT (final concentration 100 nM) also from the WA1 strain for comparison. Average IC_50_ and standard error of the means were calculated from two independent experiments with each compound concentration tested in triplicates. DMSO concentrations were always kept to ≤ 0.04% (v/v) in all wells when testing the compounds against M^pro^-E166V and to 1% (v/v) when testing the compounds against M^pro^-WT. The same concentration of DMSO was also present in control wells.

The relative fluorescence units (RFU) per second were plotted and the initial velocities were calculated, normalized to the controls (untreated protease controls wells = 0% inhibition, control wells with no substrate = 100% inhibition) and expressed as % of enzyme activity inhibition. The half maximal inhibitory concentration (IC_50_) was calculated by nonlinear regression fitting of the normalized 12-point dose response curve to the model “log(inhibitor) vs. normalized response – Variable slope” with equation: Y=100/(1+10^((LogIC_50_-X)*HillSlope)).

The data analysis was conducted in GraphPad Prism (v.9.5., GraphPad Software, La Jolla California, USA).

### 2.6 CPE-based antiviral assay

Calu-3 cells were grown in DMEM (Gibco, 41966029) supplemented with 10% FBS (Gibco, 10500064) and 1× penicillin-streptomycin (Sigma-Aldrich, PA333) and incubated at 37°C, 5% CO 2 atmosphere. The compound MP1 was tested at concentrations ranging from 20 to 0.156 *µ*M (eight 1:2 serial dilutions). MP1 was tested in two independent experiments with each concentration tested in triplicates.

One day prior to the assay, Calu-3 cells were seeded at a density of 20.000 cells/well in a 96-well plate in a final volume of 100*µ*l of cell media (DMEM supplemented with 2% FBS, 1× penicillin−streptomycin, from now on referred to as DMEM-2). After overnight incubation (37°C, 5% CO_2_ atmosphere), cells were pretreated for two hours with 100*µ*l of DMEM-2 with CP-100356 (MedChemExpress, HY-108347) added at a final concentration of 4*µ*M. After two hours, the cell media containing CP-100356 was discarded, cells were washed with 100*µ*l of PBS and infected with 200 plaque forming unites of SARS-CoV-2 (Isolate from Sweden (35)) corresponding to a MOI ∼ 0.01. After one hour the cell media with the virus was discarded, cells were washed with 100*µ*l of PBS and treated by adding 100*µ*l of DMEM-2 containing different concentrations of MP1 and CP-100356 at a final concentration of 4*µ*M. After 48h, the cell media supplemented with MP1 and CP-100356 was substituted with 100*µ*l of fresh DMEM-2 to which 10 *µ*l of a 5 mg/mL MTT (Sigma-Aldrich, M2128) solution in PBS was added. Following 4h of incubation, formazan crystals were solubilized overnight by adding 100 *µ*l of a 10% SDS, 0.01 M HCl solution. Optical density (OD) at 570 and 690 nm was read using a Tecan infinite M200 PRO plate reader (Tecan Trading AG, Switzerland). Throughout the assay, cell controls (not infected cells treated or not treated with MP1 at different concentrations) and infection controls (wells not treated with MP1 at different concentrations) were also treated with CP-100356 (4*µ*M) and DMSO concentration was kept constant to 0.25% (v/v) in all wells. OD readings at different wavelengths were subtracted, the resulting values were normalized to the controls, and EC_50_ were determined by nonlinear regression analysis using GraphPad Prism (vr.9.5, GraphPad Software, La Jolla California, USA).

### 2.7 Virus yield reduction assay

Caco-2 cells were grown in DMEM (Gibco, 41966029) supplemented with 10% FBS (Gibco, 10500064) and 1× penicillin-streptomycin (Sigma-Aldrich, PA333) and incubated at 37°C, 5% CO_2_ atmosphere. The compound MP1 was tested at 5, 0.5 and 0.05 *µ*M (eight 1:2 serial dilutions). MP1 was tested in two independent experiments with each concentration tested in triplicates.

One day prior to the assay, Caco-2 cells were seeded at a density of 20.000 cells/well in a 96-well plate in a final volume of 100*µ*l of cell media (DMEM supplemented with 2% FBS, 1× penicillin−streptomycin). After overnight incubation (37°C, 5% CO_2_ atmosphere), cells were infected with 200 plaque forming unites of SARS-CoV-2 (Isolate from Sweden(35) corresponding to a MOI ∼ 0.01. After one hour the cell media with the virus was discarded, cells were washed with 100 *µ*l of PBS and treated by adding 100 *µ*l of cell media (DMEM supplemented with 2% FBS, 1× penicillin−streptomycin) containing MP1 at different concentrations. After 48h, supernatants were collected (100 *µ*l) and mixed with 300 *µ*l of TRIzol LS (Invitrogen, ThermoFisher Scientific, Waltham, MA). The viral RNA was extracted using the Direct-zol-96 RNA Kit (Zymo Research, Irvine, CA) according to the manufacturer’s protocol. The extracted viral RNA was quantified by RT-qPCR using primers (Thermo Fisher Scientific, Waltham, MA) previously described(36) and the SuperScript III OneStep RT-PCR System with Platinum Taq DNA Polymerase kit (Invitrogen, Thermo Fisher Scientific, Waltham, MA). Target E: forward primer 5′-ACAGGTACGTTAATAGTTAATAGCGT-3′; reverse primer 5′ GTGTGCGTACTGCTGCAATAT-3′; and the probe 5′-FAM-CACTAGCCATCCTTACTGCGCTTCG-TAMRA-3′. The reaction mixture contained 12.5 *µ*l of reaction buffer (a buffer containing 0.4 mM of each dNTP, 3.2 mM Mg_2_SO_4_), 0.5 *µ*l of SuperScript III RT/Platinum Taq Mix, 0.5 *µ*l of each primer (10 μM stock concentrations), 0.25 *µ*l probe (10 μM stock concentration), 2.4 *µ*l of 25 mM magnesium sulfate, 3.35 *µ*l of nuclease-free water, and 5 *µ*l of RNA template. The RT-qPCR assay was performed on a CFX96 Touch Real-Time PCR Detection System (Bio-Rad Laboratories, Hercules CA) under the following conditions: reverse transcription at 55 °C for 30 min and 95 °C for 3 min, followed by 45 cycles of denaturation at 95 °C for 15 s, extension at 57 °C for 30 s, and collecting the fluorescence signal at 68 °C for 30s.

All samples were run in triplicate. The corresponding number of copies for each Ct was calculated from a standard curve prepared with synthetic DNA gene fragments (gBLOCKs; IDT, San Jose, CA) with a five-base-pair deletion in the amplified regions of the viral genome diluted in deionized, nuclease-free water to concentrations of 103–105 copies per *µ*l. The five-base-pairs were deleted to be able to distinguish between viral RNA and gBLOCKs during sequencing. The LODs for both genes were 101 copies per *µ*l. The relative fluorescence unit (RFU) data were obtained from the CFX Maestro Software (Bio-Rad CFX Maestro for Mac 1.1 Version 4.1.2434.0214, Bio-Rad Laboratories, Hercules, CA). Quantified viral RNA from infected wells treated with different concentrations of the compounds were normalized to the controls using GraphPad Prism (vr.9.5, GraphPad Software, La Jolla CA).

### 2.8 Compounds synthesis

All compounds were synthesized by solid-phase peptide synthesis (SPPS) on 2-chlorotrityl chloride resin (2CTC) using different Fmoc-protected natural and unnatural amino acids. All compounds prepared were either dipeptides or tripeptides terminated by a carboxylic acid as capping agent at the N-terminus. Below follows, a general synthetic procedure describing the preparation of a carboxylic acid capped tripeptide. The syntheses of the corresponding capped dipeptides were carried out in an analogous fashion using less amino acid coupling.

To a 3 mL syringe with a frit was added 2CTC resin (63 mg, 0.1 mmol, 1 equiv.), after which a solution of Fmoc-protected amino acid #1 (0.2 mmol, 2 equiv.) and *N*,*N*-diisopropylethylamine (DIPEA, 0.05 ml, 0.30 mmol, 3 equiv.) in dichloromethane (DCM, 1 ml) was prepared, and subsequently aspirated into the syringe. The resulting mixture was agitated at room temperature for 2 h to allow for coupling of amino acid #1 to the resin, after which the reaction solution was ejected and the 2CTC resin was washed with dimethyl formamide (DMF, 2×1 ml) under agitation for 30 seconds. To deactivate the remaining 2CTC functionalities on the resin, a solution of DCM/methanol/DIPEA (ratio 85:15:5, 1.05 mL) was aspirated into the syringe followed by agitation for an additional 15 min. The 2CTC resin was then washed with DMF (2×1 ml) and DCM (3×1 ml), before being subjected to subsequent Fmoc deprotection. Removal of the Fmoc group was done by treating the 2CTC resin with piperidine in methanol (80%, 1 ml) for 20 min under agitation at room temperature. The reaction solution was then ejected from the syringe, and the 2CTC resin was carefully washed using DMF (3×1 ml), methanol (2×1 ml), DCM (2×1 ml) and DMF (2×1 ml) before coupling of the next amino acid. The following method was used for coupling of amino acid #2 (0.15 mmol), and was subsequently repeated for the coupling of amino acid #3 (0.15 mmol) and the capping carboxylic acid (0.40 mmol with 3.8 equiv. HATU). The 2CTC-resin was treated in a 3 mL syringe with a frit with a solution of amino acid #2 (0.15 mmol) and DIPEA (0.07 mL, 0.4 mmol, 4 equiv.) in DMF (0.6 ml). To this mixture, was then aspirated a solution of HATU (72 mg, 0.19 mmol, 1.9 equiv.) in DMF (0.4 ml), and the corresponding solution was agitated at room temperature for 45 min, after which it was washed carefully with DMF (3×1 ml), methanol (2×1 ml), DCM (2×1 ml) and DMF (2×1 ml). Following coupling of amino acids #2 and #3, removal of the Fmoc group was done by treating the 2CTC resin with piperidine in methanol (80%, 1 ml) for 20 min under agitation at room temperature. The reaction solution was then ejected from the syringe, and the 2CTC resin was carefully washed using DMF (3×1 ml), methanol (2×1 ml), DCM (2×1 ml) and DMF (2×1 ml) before coupling of the next amino acid (or carboxylic acid in the last reaction). After completed synthesis, the target peptides were cleaved from the 2CTC resin using a mixture of hexafluoroisopropanol/DCM (700 *µ*l/300 *µ*l), followed by agitation for 10 minutes. The final product (purity >95%) was isolated after preparative HPLC purification using mobile phase: 20-60% MeCN in H_2_O with 0.05% formic acid over 8 CV with a flow of 30 mL/min.

### 2.9 Synthesis of the Dabcyl-KTSAVLQSGFRKME-EDANS substrate

EDANS NovaTag™ resin (200 mg, 0.6 mmol/g) was pre-swelled in DMF. Fmoc-Glu-OH (3 eq.), HATU (3 eq.) and DIPEA (6 eq.) in DMF was thereafter added to the resin. The reaction mixture was gently agitated in an overhead shaker for 60 min. The resin was washed several times with DMF whereafter the Fmoc protection was removed with 4% DBU in DMF followed by washings with DMF, isopropanol, methanol, H2O and DCM. The coupling procedure was repeated for the next coming Fmoc-amino acids in the following order: Fmoc-Met, Fmoc-Lys, Fmoc-Phe, Fmoc-Gly, Fmoc-Ser, Fmoc-Asn, Fmoc-Leu, Fmoc-Val, Fmoc-Ala, Fmoc-Ser, Foc-Thr and Fmoc Lys. In the final step Dabcyl was introduced by adding Dabcyl acid (3 eq.), HATU (3 eq.) and DIPEA (6 eq.) in DMF followed by agitation for 60 min. Final washing was done using DMF, iPrOH, MeOH, H2O and DCM. The final labeled peptide was cleaved from the resin using TFA:TES 95/5. The filtrate was collected and the crude product isolated by evaporation. The final product (purity 99%) was isolated after preparative HPLC purification using mobile phase: 0.1% TFA acetonitrile and 0.1% TFA in water (15– 45%).

## 3 results

### 3.1 DNA-encoded-chemical library screening

The *in vitro* affinity selection of binders from DELopen library (WuXi AppTech) was performed against recombinant avi-tagged SARS-CoV-2 M^pro^ coupled to paramagnetic beads coated with streptavidin (Figure 1). The efficiency of M^pro^ coupling to the paramagnetic beads and the activity of M^pro^ bound to the magnetics beads was tested prior to the selection experiments, ensuring that the protease was stably bound to the beads while retaining its active conformation (Supplementary Figure 1 and 2). The selection of binders was performed with or without addition of SARS-CoV-2 M^pro^ inhibitors X77 (20 *µ*M) or GC376 (5 *µ*M) that are known to bind and block the active site of M^pro^. Therefore, compounds selected in the presence of X77 or GC376 were regarded as possibly binding to an allosteric pocket. Selection was also performed against empty beads to filter out promiscuous binders of no interest. After three rounds of selection, four highly enriched binders of which one (SLL11) was a possible allosteric inhibitor, were identified by next generation sequencing (NGS) and selected for off-DNA synthesis (NGS and synthesis of compounds were provided by WuXi Apptec). The inhibitory activity of the synthesized compounds against recombinant SARS-CoV-2 M^pro^ (Table 1) was confirmed using a FRET based enzymatic assay. The inhibitory activities of the synthesized compounds against recombinant SARS-CoV-2 M^pro^ (Table 1) were confirmed using a FRET based enzymatic assay. Three out of the four synthetized compounds were active with sub micromolar IC_50_ ranging from 30 to 140 nM. The two most enriched binders against M^pro^, SLL11 and SLL12, were also the most potent (SLL11 IC_50_ = 30 nM, SLL12 IC_50_ = 53 nM) while the least enriched binder (SLL08) was not active at tested concentrations (IC_50_ > 50 *µ*M). The active compounds displayed high selectivity for SARS-CoV-2 M^pro^ and had no off-target activity against human Cathepsin S when tested up to a concentration of 50 *µ*M (Supplementary Figure 3). The two most potent compounds were synthesized and co-crystallized with M^pro^ to determine their binding pose and binding site. SLL11 and SLL12 are peptide-like compounds (Figure 2) composed of four non-natural amino acids of which P1’ present an electrophilic group. Both compounds were found to bind to the M^pro^ active site with similar binding poses proving that SLL11 was not an allosteric inhibitor despite being selected and enriched also in presence of the M^pro^ inhibitors X77 and GC376. This was not surprising as SLL11 and SLL12 shared three out of four groups.

**Table 1:** List of compounds selected from DECL libraries and relative in vitro activity.

| Compound | Enrichment score<sup>a</sup> |  |  |  | IC<sub>50</sub> (μM) |  |
| --- | --- | --- | --- | --- | --- | --- |
|  | M<sup>pro</sup> | M<sup>pro</sup><br>+X77 | M<sup>pro</sup><br>+GC376 | Beads<br>only | M<sup>pro</sup> | Cathepsin S |
| SLL07 | 135654 | 0 | 0 | 0 | 0.14 | > 50 |
| SLL08 | 11444 | 0 | 676 | 0 | > 50 | > 50 |
| SLL11 | 515485 | 20997 | 206982 | 0 | 0.03 | > 50 |
| SLL12 | 488354 | 0 | 0 | 0 | 0.053 | > 50 |

SLL07
SLL11
SLL08
SLL12
<sup>a</sup>The enrichment score quantify how abundant a compound was after selection, e.g. a compound with enrichment score = 100 was 100 times more abundant than average

**Figure 1:**
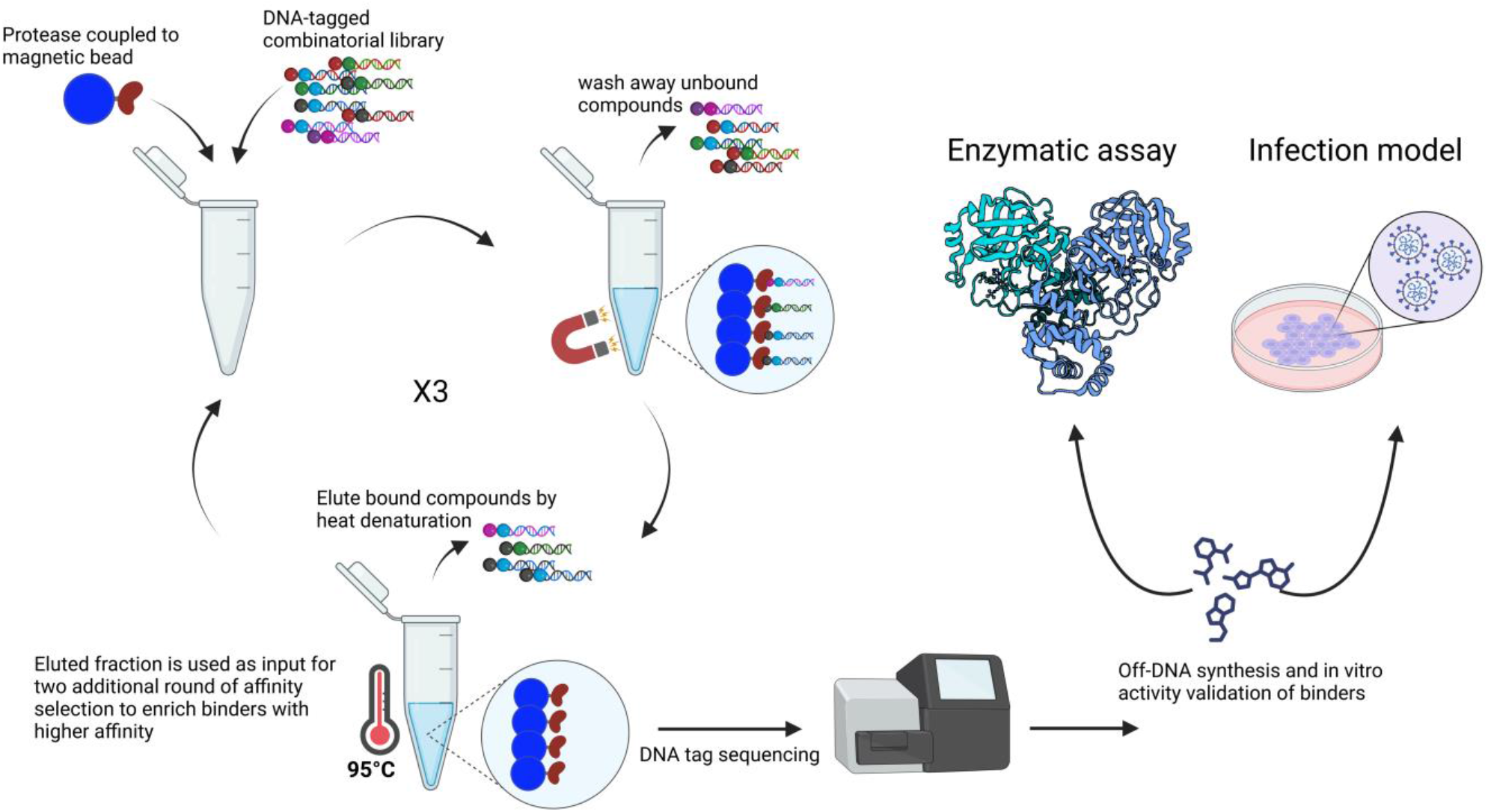
Workflow of hit identification and validation using DNA-encoded chemical libraries. Novel inhibitors of SARS-CoV-2 M^pro^ were identified from the DELopen library containing 4.2 billion DNA-tagged compounds. After three rounds of affinity selection, the unique DNA-tag of binders was sequenced allowing the identification of highly enriched compounds with their numeric building block codes. The molecular structures of the most promising compounds were subsequentially disclosed by WuXi App Tec, four compounds were synthetized off-DNA and tested to confirm their in vitro inhibitory activity against M^pro^ and antiviral activity against SARS-CoV-2 in cell-based assays.

**Figure 2:**
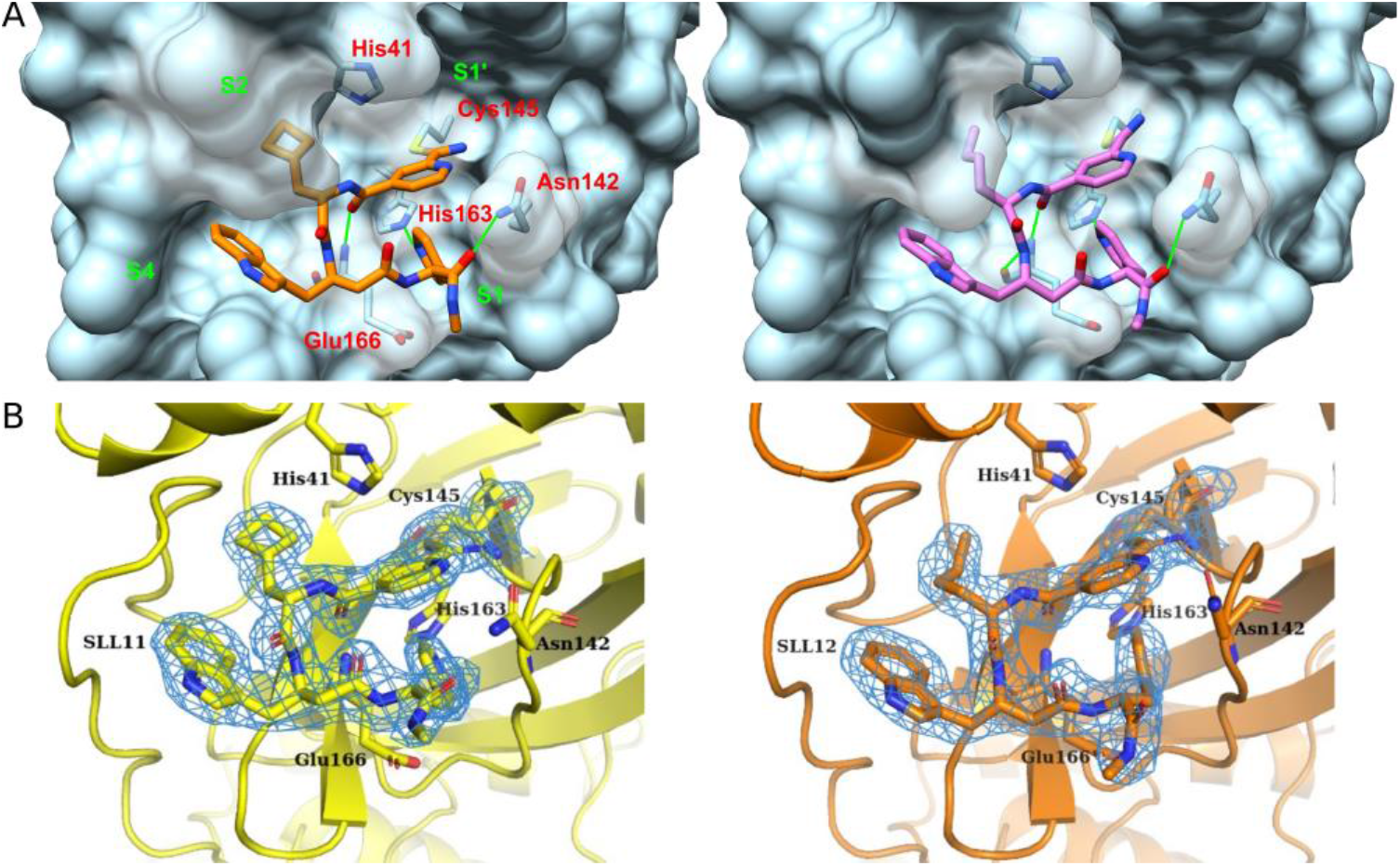
Compound’s general structure and interactions made with SARS-CoV-2 M^pro^. (A) Crystal structures of compounds SLL11 (Provisory PDB ID 9EO6), shown in orange, and SLL12 (Provisory PDB ID 9EOR), shown in purple bound to SARS-CoV-2 M^pro^ active site. The two residues forming the enzyme’s catalytic dyad (His41 and Cys145) and the residues found interacting with SLL11 and SLL12 are shown in red. The subsites of M^pro^ active site and predicted hydrogen bonds are labeled and drawn in green respectively. (B) Electron density maps (2fo-fc) covering the ligands SLL11 (yellow) and SLL12 (orange) contoured at 1.2 sigma level.

The compounds interacted with M^pro^ mainly via a covalent bond formed between the electrophilic P1’ group and the catalytic Cys145 close to the S1’ subsite and a hydrogen bond formed between the P3 group and His163 in the S1 subsite. While P2 and P3 hydrophobic residues occupied the S2 and S4 subsite. Hydrogen bonds were also formed between the backbone of the compounds and Glu166 and between the C-terminal amine of SLL11 and M^pro^ Asn142 side chain.

### 3.2 Structure activity relationship study

Compound MP1 was obtained substituting the C-terminus of compound SLL11 (IC_50_ = 30 nM) with a carboxylic acid group to increase the compound solubility (Table 2). This structural difference caused no loss of activity and most likely did not affect its binding mode (Supplementary figure 4), thus compound MP1 (IC_50_ = 25 nM) was used as starting point and reference for the design of analogues and comparison of inhibitory activity (Table 2). The inhibitory activity of analogues against recombinant SARS-CoV-2 M^pro^ was first screened at a concentration of 1 *µ*M and analogues inhibiting M^pro^ activity by 80% or more were further tested to determine their IC_50_ (Dose response curves are shown in Supplementary Figure 5).

**Table 2:**
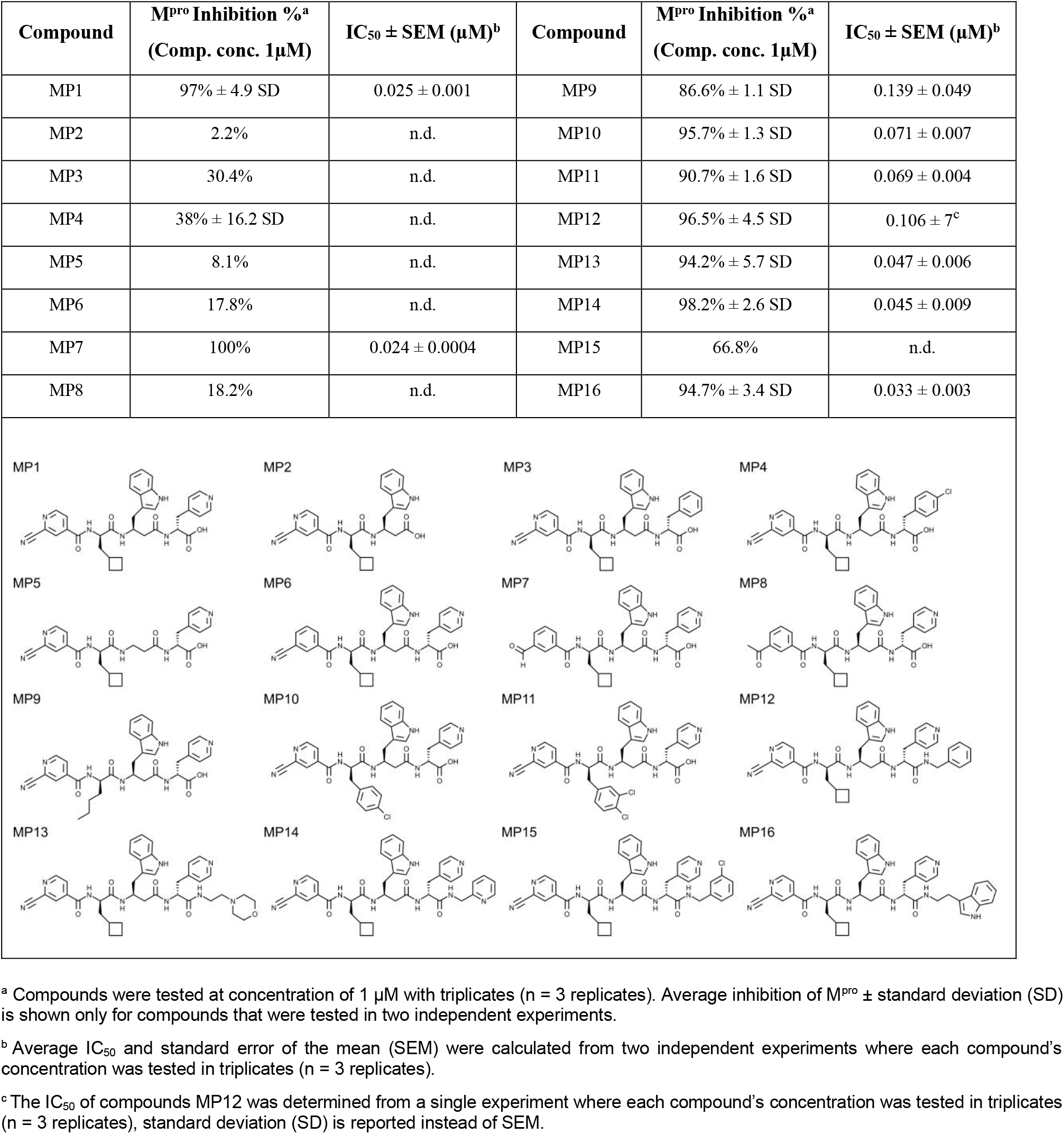
List of MP1 analogues and relative Inhibitory activity against SARS-CoV-2 M^pro^

We assumed that the covalent bond formed between the N-terminal electrophilic group and Cys145 was essential for MP1 activity and started designing analogues with modification at the C-terminus of the molecule. First, analogues presenting variations of the P3 and P2 residues, were designed to assess if the size of MP1 could be reduced while retaining activity. As observed from the crystallographic binding poses, the P3 residue occupies the S1 subsite of M^pro^ and forms a hydrogen bond with His163. Therefore, removing the P3 residues (Compound MP2) or substituting P3 with groups lacking a hydrogen bond acceptor (Compounds MP3 and MP4) were poorly tolerated, reducing the compounds inhibitory activity to ∼2% and ∼30% respectively. Removing the side group of the P2 residue while maintaining the portion of backbone connecting P3 to the P1 residue (Compound MP5) also sharply reduced the inhibitory effect to ∼8%. Since both P3 and P2 groups were essential for binding, and attempts to shorten MP1 were unsuccessful, we next assessed the contribution of the P1’ electrophilic group to the binding of MP1 to the active of SARS-CoV-2 M^pro^. As expected, substituting the P1’ residue with a less reactive analogue (Compounds MP6) was detrimental (∼18% residual inhibitory activity) while a highly reactive aldehyde (compound MP7, IC_50_ = 24 nM) was equipotent to the original electrophile group of MP1. A bulkier electrophile group at P1’ (Compounds MP8) also reduced the inhibitory activity against M^pro^ to ∼ 18%. Lastly, the analogues MP9, MP10 and MP11 were designed to evaluate variants of the P1 residues. Exchanging the hydrophobic P1 group to other hydrophobic aliphatic (MP9 IC_50_ = 139 nM) or planar aromatic groups (MP10 IC_50_ = 71 nM and MP11 IC_50_ = 69 nM) reduced the IC_50_ of the analogues up to 4-fold. However, all the analogues maintained a sub micromolar activity with IC_50_ in the low nanomolar range.

Overall, all residues constituting MP1 seemed to be well optimized, so we next explored the post-modification of the C-terminus with various amide groups. Analogues with polar amide substituents (compounds MP13 IC_50_ = 47 nM, MP14 IC_50_ = 45 nM and MP16 IC_50_ = 47 nM) were ∼2 fold more potent than analogues with a non-polar amide groups (MP12 IC_50_ = 106 nm), suggesting that this substituent might take part in hydrogen bonding with residues located near the S1 subsite of M^pro^.

### 3.3 Antiviral activity of MP1 against SARS-CoV-2 in cell-based assays

MP1 was chosen to test the antiviral activity of the novel scaffold discovered in this study. The compound cell permeability was first evaluated Caco-2 cells (human colon epithelial cells) that were cultured on a permeable filter. MP1 showed good apical to basolateral (A-B) permeability (*P*app A-B = 5.4 x 10^6^ ± 3.6 x 10^6^ cm/s) and lower basolateral to apical (B-A) permeability (*P*app B-A = 2.5 x 10^6^ ± 0.6 x 10^7^ cm/s) with no observable efflux of the compound (*P*app B-A / *P*app A-B = 0.5). Accordingly, MP1 was active in Caco-2 cells and showed dose response inhibition of the viral replication measured by RT-qPCR (Figure 3A). Since Caco-2 cells are highly permissive for SARS-CoV-2 infection but do not develop CPE, we tested the capacity of MP1 to inhibit CPE development and increase cell viability in infected Calu-3 cells (Human lung epithelial cells). Unexpectedly, MP1 had no protective effect in infected Calu-3 at concentrations as high as 20 *µ*M (data not shown). Based on previous studies reporting the P-glycoprotein (P-gp) mediated efflux of M^pro^ peptide-like inhibitors (12, 37, 38), we assumed MP1 being a substrate of P-gp. When tested in combination with the P-gp inhibitor CP-100356 (4 *µ*M), MP1 showed dose dependent inhibition of CPE development with EC_50_ = 2.3 ± 1.1 *µ*M and observable cytotoxic effect (Fig. 3B).

**Figure 3:**
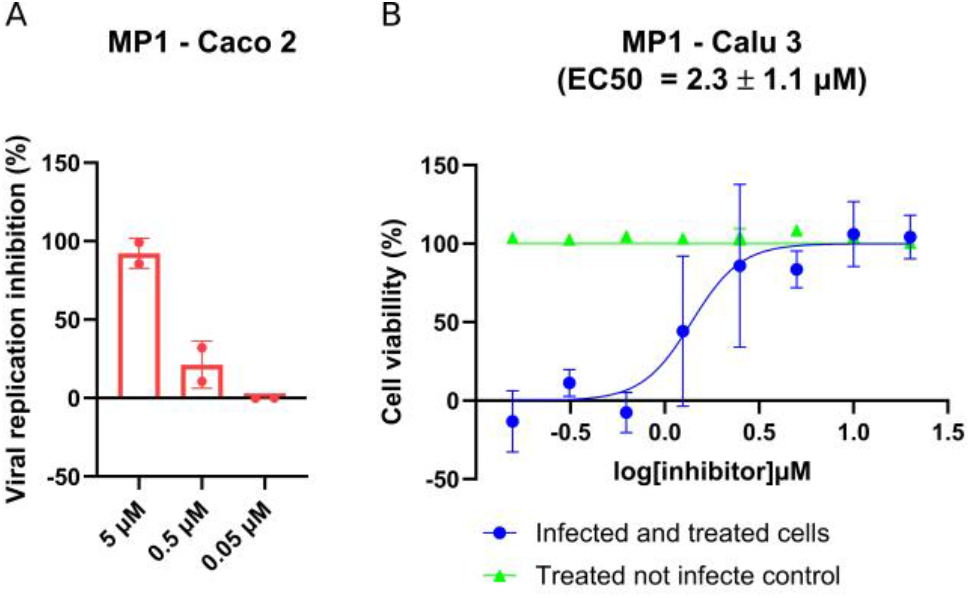
Antiviral effect of MP1 against SARS-CoV-2. (A) The Inhibitory effect on viral replication was assessed using a yield reduction assay. Caco-2 cells were infected with SARS-CoV-2 (MOI 0.01) and treated with 5, 0.5 or 0.05 *µ*M of MP1. Supernatants were collected 48h post infection and viral copy number per ml were quantified by RT-qPCR. The reduction in viral copy number per ml from two independent experiments is reported as inhibition of viral replication (%) ± SD. (B) Inhibitory effect on CPE development induced by SARS-CoV-2 infection. Calu-3 cells were infected (MOI 0.01) and treated with different concentrations of MP1, 48h post infection cell viability was assessed by MTT assay. The average EC_50_ from two independent experiments is reported ± the standard error of the mean.

### 3.4 Effect of the E166V M^pro^ variant on MP1 and MP7 inhibitory activity

The inhibitory activity of compounds MP1 and MP7 was tested against recombinant M^pro^ carrying the E166V (M^pro^-E166V) variant, known to confer resistance against nirmatrelvir. Compounds MP1, MP7 and nirmatrelvir, used as a reference, had comparable IC_50_ in the low nanomolar range against the recombinant wild type M^pro^ (M^pro^-WT). The inhibitory activity of the three compounds was significantly decreased to 36.6 *µ*M (MP1), 21.1 *µ*M (MP7) and 15.7 *µ*M (nirmatrelvir) and against M^pro^-E166V (Figure 4).

**Figure 4:**
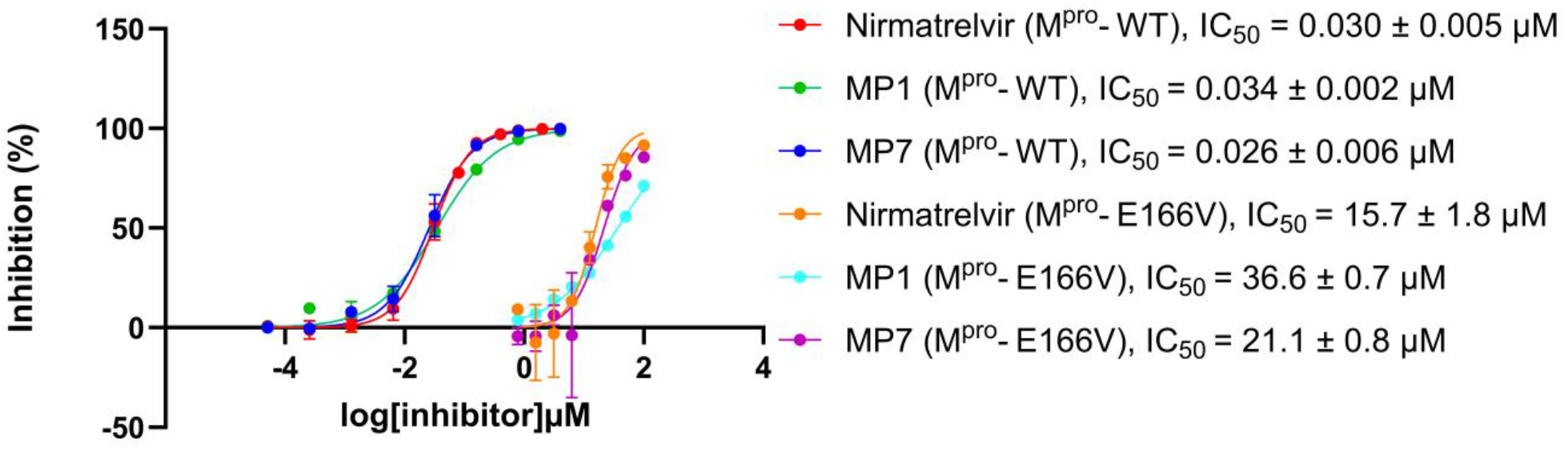
Effect of the E166V variant on the inhibitory activity of MP1 and MP7 compounds. Dose response curves of MP1 and MP7 tested against recombinants wild type (M^pro^-WT) and mutated M^pro^ carrying the E166V (M^pro^-E166V). IC_50_ values are reported as average ± SEM from two independent experiments perfumed with triplicates. Nirmatrelvir was also tested for comparison.

## 4 Discussion

In this study we report the identification of novel inhibitors of SARS-COV-2 M^pro^ from the commercially available “DELopen” DNA encoded chemical library (DECL).

Three potent compounds (SLL07, SLL11 and SLL12) with IC_50_ in the low nanomolar range were directly identified through the cost-efficient affinity screening of 4.2 billion combinatorial molecules. The selected peptide-like compounds were formed by three unnatural amino acids capped at the N-terminus by a carboxylic acid derivative carrying a reactive electrophilic group. Crystallographic studies of SLL11 and SLL12 binding poses showed that side groups of all four residues (P1’, P1, P2 and P3) occupied one of the major subsites of M^pro^ active site (S1’, S1, S2 and S4) with a novel arrangement of the peptide-like backbone to fit the binding site. The backbone of compounds SLL11 and SLL12 in the crystal structures assumed an unprecedented semi-cyclic conformation with the N-terminus and C-terminus stacked and possibly forming a T-shaped non-bonded π-π interaction. (Figure 2). In this conformation the N-terminal electrophilic group binds Cys145 close to the S1’ pocket while the C-terminal pyridine group binds in the S1 pocket. This C-terminus to N-terminus orientation is opposite to the orientation of the natural substrate of M^pro^(39) and its substrate-derived inhibitors. This results in the peptide backbone of SLL11 and SLL12 assuming the typical conformation of a D-peptide with the amino groups presented on the left side of the backbone alpha carbon as opposed to conventional L-amino acid backbones displaying them on the right side. This unusual confirmation becomes more evident when the binding poses of SLL11 and nirmatrelvir bound to the M^pro^ of SARS-CoV-2 are compared as shown in the Supplementary Figure 6.

Attempts to further improve the compounds’ potency by systematically varying one of the building blocks residues resulted in the design of several analogues with IC_50_ lower than 100 nM. However, SLL11 (and its close analogue MP1) remained the most potent inhibitor (IC_50_ = ≤ 30 nM) proving that the screening of large combinatorial libraries can identify well optimized compounds without the need for several iterative cycles of medicinal chemistry.

Reducing the molecular weight thus improving the physiochemical properties of the compounds proved to be difficult. The inhibitory activity of analogues with a less reactive electrophile at P1’ was almost completely abrogated, suggesting that a covalent interaction with M^pro^ was essential. Co-crystallization of SLL11 and SLL12 with SARS-CoV-2 M^pro^ confirmed that the N-terminal nitrile electrophilic group formed a covalent bond with Cys145 of enzyme’s catalytic dyad as reported for several other peptide-like inhibitors of M^pro^ (12, 38, 40, 41). Likewise, removing the whole P2 residue or P3 side group was not possible without causing a drop in compounds inhibitory activity. P3 formed an essential hydrogen bond with S2 His163 that mimics the interaction with the glutamine residue of its natural substrate (42). The side group of the P2 residue did not form any hydrogen bond with residues in the S4 pocket but was still essential for activity, suggesting that P2 contributes with more than just connecting P3 to the remaining part of the molecule.

When tested in cell-based assays, MP1 showed antiviral activity against SARS-CoV-2 in the low micromolar range in both Caco-2 and Calu-3 (EC_50_ = 2.3 ± 1.1 *µ*M), two cell lines commonly used as infection model for SARS-CoV-2. However, while no efflux was observed in Caco-2 permeability test, the addition of CP-100356 was necessary to prevent MP1 efflux mediated by P-glycoprotein (P-gp) in Calu-3 cells. This finding supports a previous report of Calu-3 cells expressing P-gp at a higher level than Caco-2 cells (43). The relatively low activity of MP1 in cell-based assay could be explained by the compound’s sensitivity to proteolytic degradation as both SLL11 and SLL12 showed low metabolic stability in human and mouse microsomes (supplementary Table 2). Cyclization or substitution of selected residues with D-enantiomers are two strategies that could improve the metabolic stability of the peptide-like compounds reported in this study and will be subject of further studies in our lab.

Finally, the in vitro inhibitory activity of MP1 and MP7 was evaluated against recombinant M^pro^ carrying the variant E166V that cause high fold resistance to nirmatrelvir (23, 44, 45). Nirmatrelvir had a 523-fold resistance when tested against the M^pro^-E166V in our FRET based enzymatic assay. The observed increase in nirmatrelvir’s IC_50_ against SARS-CoV-2 M^pro^ was comparable to the 473-fold increase reported by Lan et al. in a similar enzymatic assay (23). The compounds MP1 and MP7 were also subject to decreased inhibitory activity and had 1076 and 811-fold resistance respectively when tested against M^pro^-E166V. The dramatic effect of E166V mutation on nirmatrelvir activity was shown to be due to steric clashes in S1 preventing the formation of hydrogen bonds between nirmatrelvir’s γ-lactam ring, residues His163 and possibly also Asp166 itself or Phe140. As a result of the disrupted key interaction in S1, the nitrile warhead is moved out of position making it more difficult to interact with Cys145 (23, 45).

MP1 and MP7 also interact with residue His163 trough the hydrogen bonding and Cys145 with their electrophilic warheads at P1’. It is therefore plausible that similar dynamics are the cause of the reduced activity observed *in vitro* for compounds MP1 and MP7.

In conclusion, the affinity-based selection of a ultralarge DECL resulted in the direct identification of novel peptide-like inhibitors of SARS-CoV-2 M^pro^ with *in vitro* inhibitory activity in the low nanomolar range against recombinant M^pro^ and inhibitory activity in the low micromolar range in cell-based assays. As demonstrated herein, the use of this technique allows the rapid exploration of large chemical spaces and could greatly expedite the discovery and optimization of novel active compounds against SARS-CoV-2 M^pro^.

## CRediT author statement

**Dario Akaberi:** Conceptualization, Methodology, Validation, Formal analysis, Investigation, Writing - Original Draft, Writing - Review & Editing, Visualization. **Monireh Pourghasemi Lati:** Methodology, Validation, Investigation. **Janina Krambrich:** Investigation. **Julia Berger:** Validation. **Grace Neilsen:** Methodology. **Emilia Strandback:** Methodology. **Pauliina Turunen:** Validation, Investigation. **Johan Wannberg:** Methodology, Validation. **Hjalmar Gullberg:** Validation, Investigation. **Martin Moche:** Methodology, Formal analysis, Investigation, Visualization. **Praveen Kumar Chinthakindi:** Methodology. **Tomas Nyman:** Supervision. **Stefan G. Sarafianos:** Supervision. **Anja Sandström:** Supervision. **Josef Järhult:** Supervision. **Kristian Sandberg:** Resources, Supervision. **Åke Lundkvist:** Resources, Supervision. **Oscar Verho:** Methodology, Formal analysis, Resources, Writing - Review & Editing, Supervision. **Johan Lennerstrand:** Conceptualization, Resources, Writing - Review & Editing, Supervision, Project administration, Funding acquisition.

## Acknowledgments

We would like to thank the Protein Science Facility (PSF) team at the Karolinska Institutet, for the production of recombinant SARS-CoV-2 main protease used in the enzymatic assays and, Martin Walsh (Diamond Light Source, UK) for the production of the recombinant Avi-tagged main protease used for the affinity screening of DECL. The 3D structures of proteins were determined at PSF, we thank MAX IV light source for allocation of BioMAX beamtime to project 20200071 and Diamond Light Source for access to i04 via MX21625. The calculation of 3D structures was performed at NSC Tetralith cluster provided by the Swedish National Infrastructure for Computing (SNIC) and software installed by the PReSTO project funded by the Swedish Research Council through grant agreement no. 2018-05973 (SNIC) and 2018-06479 (PReSTO). The DELopen library and the off-DNA synthesis of hits were provided by WuXi App Tec. We also thank Pawel Baranczewski for performing metabolic stability studies and Richard Svensson for performing cell permeability studies in Caco-2 cell lines.

## References

1. Viana R, Moyo S, Amoako DG, Tegally H, Scheepers C, Althaus CL, Anyaneji UJ, Bester PA, Boni MF, Chand M, Choga WT, Colquhoun R, Davids M, Deforche K, Doolabh D, du Plessis L, Engelbrecht S, Everatt J, Giandhari J, Giovanetti M, Hardie D, Hill V, Hsiao N-Y, Iranzadeh A, Ismail A, Joseph C, Joseph R, Koopile L, Kosakovsky Pond SL, Kraemer MUG, Kuate-Lere L, Laguda-Akingba O, Lesetedi-Mafoko O, Lessells RJ, Lockman S, Lucaci AG, Maharaj A, Mahlangu B, Maponga T, Mahlakwane K, Makatini Z, Marais G, Maruapula D, Masupu K, Matshaba M, Mayaphi S, Mbhele N, Mbulawa MB, Mendes A, Mlisana K, Mnguni A, Mohale T, Moir M, Moruisi K, Mosepele M, Motsatsi G, Motswaledi MS, Mphoyakgosi T, Msomi N, Mwangi PN, Naidoo Y, Ntuli N, Nyaga M, Olubayo L, Pillay S, Radibe B, Ramphal Y, Ramphal U, San JE, Scott L, Shapiro R, Singh L, Smith-Lawrence P, Stevens W, Strydom A, Subramoney K, Tebeila N, Tshiabuila D, Tsui J, van Wyk S, Weaver S, Wibmer CK, Wilkinson E, Wolter N, Zarebski AE, Zuze B, Goedhals D, Preiser W, Treurnicht F, Venter M, Williamson C, Pybus OG, Bhiman J, Glass A, Martin DP, Rambaut A, Gaseitsiwe S, von Gottberg A, de Oliveira T. 2022. Rapid epidemic expansion of the SARS-CoV-2 Omicron variant in southern Africa. 7902. Nature 603:679–686.

2. Kurhade C, Zou J, Xia H, Liu M, Chang HC, Ren P, Xie X, Shi P-Y. 2022. Low neutralization of SARS-CoV-2 Omicron BA.2.75.2, BQ.1.1 and XBB.1 by parental mRNA vaccine or a BA.5 bivalent booster. Nat Med 1–4.

3. Lyke KE, Atmar RL, Islas CD, Posavad CM, Szydlo D, Paul Chourdhury R, Deming ME, Eaton A, Jackson LA, Branche AR, El Sahly HM, Rostad CA, Martin JM, Johnston C, Rupp RE, Mulligan MJ, Brady RC, Frenck RW, Bäcker M, Kottkamp AC, Babu TM, Rajakumar K, Edupuganti S, Dobrzynski D, Coler RN, Archer JI, Crandon S, Zemanek JA, Brown ER, Neuzil KM, Stephens DS, Post DJ, Nayak SU, Suthar MS, Roberts PC, Beigel JH, Montefiori DC, Husson JS, Price A, Whitaker JA, Keitel WA, Falsey AR, Shannon I, Graciaa D, Rouphael N, Anderson EJ, Kamidani S, Muniz GB, Bhatnagar S, Wald A, Berman M, Porterfield L, Stanford A, Dong JL, Carsons SE, Badillo D, Parker S, Dickey M, Larsen SE, Hural J, Ingersoll B, Lee M, Lai L, Floyd K, Ellis M, Moore KM, Manning K, Foster SL, Patel M. 2022. Rapid decline in vaccine-boosted neutralizing antibodies against SARS-CoV-2 Omicron variant. Cell Reports Medicine 3:100679.

4. Wang Q, Iketani S, Li Z, Liu L, Guo Y, Huang Y, Bowen AD, Liu M, Wang M, Yu J, Valdez R, Lauring AS, Sheng Z, Wang HH, Gordon A, Liu L, Ho DD. 2022. Alarming antibody evasion properties of rising SARS-CoV-2 BQ and XBB subvariants. Cell 0.

5. Keam SJ. 2022. Tixagevimab + Cilgavimab: First Approval. Drugs 82:1001–1010.

6. Kim JY, Săndulescu O, Preotescu L-L, Rivera-Martínez NE, Dobryanska M, Birlutiu V, Miftode EG, Gaibu N, Caliman-Sturdza O, Florescu S-A, Shi HJ, Streinu-Cercel A, Streinu-Cercel A, Lee SJ, Kim SH, Chang I, Bae YJ, Suh JH, Chung DR, Kim SJ, Kim MR, Lee SG, Park G, Eom JS. 2022. A Randomized Clinical Trial of Regdanvimab in High-Risk Patients With Mild-to-Moderate Coronavirus Disease 2019. Open Forum Infect Dis 9:ofac406.

7. Lee S, Lee SO, Lee JE, Kim K-H, Lee SH, Hwang S, Kim S-W, Chang H-H, Kim Y, Bae S, Kim A-S, Kwon KT. 2022. Regdanvimab in patients with mild-to-moderate SARS-CoV-2 infection: A propensity score-matched retrospective cohort study. Int Immunopharmacol 106:108570.

8. Zhang L, Lin D, Sun X, Curth U, Drosten C, Sauerhering L, Becker S, Rox K, Hilgenfeld R. 2020. Crystal structure of SARS-CoV-2 main protease provides a basis for design of improved α-ketoamide inhibitors. Science 368:409–412.

9. V’kovski P, Kratzel A, Steiner S, Stalder H, Thiel V. 2021. Coronavirus biology and replication: implications for SARS-CoV-2. 3. Nat Rev Microbiol 19:155–170.

10. Roe MK, Junod NA, Young AR, Beachboard DC, Stobart CC. 2021. Targeting novel structural and functional features of coronavirus protease nsp5 (3CLpro, Mpro) in the age of COVID-19. J Gen Virol 102:001558.

11. Luttens A, Gullberg H, Abdurakhmanov E, Vo DD, Akaberi D, Talibov VO, Nekhotiaeva N, Vangeel L, De Jonghe S, Jochmans D, Krambrich J, Tas A, Lundgren B, Gravenfors Y, Craig AJ, Atilaw Y, Sandström A, Moodie LWK, Lundkvist Å, van Hemert MJ, Neyts J, Lennerstrand J, Kihlberg J, Sandberg K, Danielson UH, Carlsson J. 2022. Ultralarge Virtual Screening Identifies SARS-CoV-2 Main Protease Inhibitors with Broad-Spectrum Activity against Coronaviruses. J Am Chem Soc 144:2905–2920.

12. Owen DR, Allerton CMN, Anderson AS, Aschenbrenner L, Avery M, Berritt S, Boras B, Cardin RD, Carlo A, Coffman KJ, Dantonio A, Di L, Eng H, Ferre R, Gajiwala KS, Gibson SA, Greasley SE, Hurst BL, Kadar EP, Kalgutkar AS, Lee JC, Lee J, Liu W, Mason SW, Noell S, Novak JJ, Obach RS, Ogilvie K, Patel NC, Pettersson M, Rai DK, Reese MR, Sammons MF, Sathish JG, Singh RSP, Steppan CM, Stewart AE, Tuttle JB, Updyke L, Verhoest PR, Wei L, Yang Q, Zhu Y. 2021. An oral SARS-CoV-2 Mpro inhibitor clinical candidate for the treatment of COVID-19. Science 374:1586–1593.

13. Unoh Y, Uehara S, Nakahara K, Nobori H, Yamatsu Y, Yamamoto S, Maruyama Y, Taoda Y, Kasamatsu K, Suto T, Kouki K, Nakahashi A, Kawashima S, Sanaki T, Toba S, Uemura K, Mizutare T, Ando S, Sasaki M, Orba Y, Sawa H, Sato A, Sato T, Kato T, Tachibana Y. 2022. Discovery of S-217622, a Noncovalent Oral SARS-CoV-2 3CL Protease Inhibitor Clinical Candidate for Treating COVID-19. J Med Chem 65:6499–6512.

14. Shionogi, press release. 2022. Xocova® (Ensitrelvir Fumaric Acid) Tablets 125mg Approved in Japan for the Treatment of SARS-CoV-2 Infection, under the Emergency Regulatory Approval System. https://www.shionogi.com/us/en/news/2022/11/xocova-ensitrelvir-fumaric-acid-tablets-125mg-approved-in-japan-for-the-treatment-of-sars-cov-2-infection,-under-the-emergency-regulatory-approval-system.html. Retrieved 10 January 2023.

15. Amblard F, LeCher JC, D. R, Zhou S, Liu P, Goh SL, Tao S, Patel D, Downs-Bowen J, Zandi K, Zhang H, Chaudhry G, McBrayer T, Muczynski M, Al-Homoudi A, Engel J, Lan S, Sarafianos SG, Kovari LC, Schinazi RF. 2024. Synthesis and biological evaluation of novel peptidomimetic inhibitors of the coronavirus 3C-like protease. European Journal of Medicinal Chemistry 268:116263.

16. Han SH, Goins CM, Arya T, Shin W-J, Maw J, Hooper A, Sonawane DP, Porter MR, Bannister BE, Crouch RD, Lindsey AA, Lakatos G, Martinez SR, Alvarado J, Akers WS, Wang NS, Jung JU, Macdonald JD, Stauffer SR. 2022. Structure-Based Optimization of ML300-Derived, Noncovalent Inhibitors Targeting the Severe Acute Respiratory Syndrome Coronavirus 3CL Protease (SARS-CoV-2 3CLpro). J Med Chem 65:2880–2904.

17. Huang C, Shuai H, Qiao J, Hou Y, Zeng R, Xia A, Xie L, Fang Z, Li Y, Yoon C, Huang Q, Hu B, You J, Quan B, Zhao X, Guo N, Zhang S, Ma R, Zhang J, Wang Y, Yang R, Zhang S, Nan J, Xu H, Wang F, Lei J, Chu H, Yang S. 2023. A new generation Mpro inhibitor with potent activity against SARS-CoV-2 Omicron variants. 1. Sig Transduct Target Ther 8:1–13.

18. Douangamath A, Fearon D, Gehrtz P, Krojer T, Lukacik P, Owen CD, Resnick E, Strain-Damerell C, Aimon A, Ábrányi-Balogh P, Brandão-Neto J, Carbery A, Davison G, Dias A, Downes TD, Dunnett L, Fairhead M, Firth JD, Jones SP, Keeley A, Keserü GM, Klein HF, Martin MP, Noble MEM, O’Brien P, Powell A, Reddi RN, Skyner R, Snee M, Waring MJ, Wild C, London N, von Delft F, Walsh MA. 2020. Crystallographic and electrophilic fragment screening of the SARS-CoV-2 main protease. 1. Nat Commun 11:5047.

19. Chamakuri S, Lu S, Ucisik MN, Bohren KM, Chen Y-C, Du H-C, Faver JC, Jimmidi R, Li F, Li J-Y, Nyshadham P, Palmer SS, Pollet J, Qin X, Ronca SE, Sankaran B, Sharma KL, Tan Z, Versteeg L, Yu Z, Matzuk MM, Palzkill T, Young DW. 2021. DNA-encoded chemistry technology yields expedient access to SARS-CoV-2 Mpro inhibitors. Proceedings of the National Academy of Sciences 118:e2111172118.

20. Ge R, Shen Z, Yin J, Chen W, Zhang Q, An Y, Tang D, Satz AL, Su W, Kuai L. 2022. Discovery of SARS-CoV-2 main protease covalent inhibitors from a DNA-encoded library selection. SLAS Discov 27:79–85.

21. Satz AL, Brunschweiger A, Flanagan ME, Gloger A, Hansen NJV, Kuai L, Kunig VBK, Lu X, Madsen D, Marcaurelle LA, Mulrooney C, O’Donovan G, Sakata S, Scheuermann J. 2022. DNA-encoded chemical libraries. 1. Nat Rev Methods Primers 2:1–17.

22. Akaberi D, Krambrich J, Ling J, Luni C, Hedenstierna G, Järhult JD, Lennerstrand J, Lundkvist Å. 2020. Mitigation of the replication of SARS-CoV-2 by nitric oxide in vitro. Redox Biology 37:101734.

23. Lan S, Neilsen G, Slack RL, Cantara WA, Castaner AE, Lorson ZC, Lulkin N, Zhang H, Lee J, Cilento ME, Tedbury PR, Sarafianos SG. 2023. Nirmatrelvir Resistance in SARS-CoV-2 Omicron_BA.1 and WA1 Replicons and Escape Strategies. bioRxiv 10.1101/2022.12.31.522389.

24. Ursby T, Åhnberg K, Appio R, Aurelius O, Barczyk A, Bartalesi A, Bjelčić M, Bolmsten F, Cerenius Y, Doak RB, Eguiraun M, Eriksson T, Friel RJ, Gorgisyan I, Gross A, Haghighat V, Hennies F, Jagudin E, Norsk Jensen B, Jeppsson T, Kloos M, Lidon-Simon J, de Lima GMA, Lizatovic R, Lundin M, Milan-Otero A, Milas M, Nan J, Nardella A, Rosborg A, Shilova A, Shoeman RL, Siewert F, Sondhauss P, Talibov VO, Tarawneh H, Thånell J, Thunnissen M, Unge J, Ward C, Gonzalez A, Mueller U. 2020. BioMAX -the first macromolecular crystallography beamline at MAX IV Laboratory. J Synchrotron Radiat 27:1415–1429.

25. Materlik G, Rayment T, Stuart DI. 2015. Diamond Light Source: status and perspectives. Philos Trans A Math Phys Eng Sci 373:20130161.

26. Kabsch W. 2010. XDS. Acta Crystallogr D Biol Crystallogr 66:125–132.

27. Sparta KM, Krug M, Heinemann U, Mueller U, Weiss MS. 2016. XDSAPP2.0. J Appl Cryst 49:1085–1092.

28. Winn MD, Ballard CC, Cowtan KD, Dodson EJ, Emsley P, Evans PR, Keegan RM, Krissinel EB, Leslie AGW, McCoy A, McNicholas SJ, Murshudov GN, Pannu NS, Potterton EA, Powell HR, Read RJ, Vagin A, Wilson KS. 2011. Overview of the CCP4 suite and current developments. Acta Crystallogr D Biol Crystallogr 67:235–242.

29. Evans PR, Murshudov GN. 2013. How good are my data and what is the resolution? Acta Crystallogr D Biol Crystallogr 69:1204–1214.

30. McCoy AJ, Grosse-Kunstleve RW, Adams PD, Winn MD, Storoni LC, Read RJ. 2007. Phaser crystallographic software. J Appl Cryst 40:658–674.

31. Murshudov GN, Skubák P, Lebedev AA, Pannu NS, Steiner RA, Nicholls RA, Winn MD, Long F, Vagin AA. 2011. REFMAC5 for the refinement of macromolecular crystal structures. Acta Crystallogr D Biol Crystallogr 67:355–367.

32. Emsley P, Lohkamp B, Scott WG, Cowtan K. 2010. Features and development of Coot. Acta Crystallogr D Biol Crystallogr 66:486–501.

33. Long F, Nicholls RA, Emsley P, Gražulis S, Merkys A, Vaitkus A, Murshudov GN. 2017. AceDRG: a stereochemical description generator for ligands. Acta Crystallogr D Struct Biol 73:112–122.

34. Nicholls RA, Wojdyr M, Joosten RP, Catapano L, Long F, Fischer M, Emsley P, Murshudov GN. 2021. The missing link: covalent linkages in structural models. Acta Crystallogr D Struct Biol 77:727–745.

35. Ling J, Hickman RA, Frithiof R, Hultström M, Järhult JD, Lundkvist Å, Lipcsey M. 2023. Infectious SARS-CoV-2 is rarely present in the nasopharynx samples collected from Swedish hospitalized critically ill COVID-19 patients. Ir J Med Sci 192:227–229.

36. Corman VM, Landt O, Kaiser M, Molenkamp R, Meijer A, Chu DK, Bleicker T, Brünink S, Schneider J, Schmidt ML, Mulders DG, Haagmans BL, Veer B van der, Brink S van den, Wijsman L, Goderski G, Romette J-L, Ellis J, Zambon M, Peiris M, Goossens H, Reusken C, Koopmans MP, Drosten C. 2020. Detection of 2019 novel coronavirus (2019-nCoV) by real-time RT-PCR. Eurosurveillance 25:2000045.

37. Boras B, Jones RM, Anson BJ, Arenson D, Aschenbrenner L, Bakowski MA, Beutler N, Binder J, Chen E, Eng H, Hammond H, Hammond J, Haupt RE, Hoffman R, Kadar EP, Kania R, Kimoto E, Kirkpatrick MG, Lanyon L, Lendy EK, Lillis JR, Logue J, Luthra SA, Ma C, Mason SW, McGrath ME, Noell S, Obach RS, O’ Brien MN, O’Connor R, Ogilvie K, Owen D, Pettersson M, Reese MR, Rogers TF, Rosales R, Rossulek MI, Sathish JG, Shirai N, Steppan C, Ticehurst M, Updyke LW, Weston S, Zhu Y, White KM, García-Sastre A, Wang J, Chatterjee AK, Mesecar AD, Frieman MB, Anderson AS, Allerton C. 2021. Preclinical characterization of an intravenous coronavirus 3CL protease inhibitor for the potential treatment of COVID19. Nat Commun 12:6055.

38. Hoffman RL, Kania RS, Brothers MA, Davies JF, Ferre RA, Gajiwala KS, He M, Hogan RJ, Kozminski K, Li LY, Lockner JW, Lou J, Marra MT, Mitchell LJ Jr, Murray BW, Nieman JA, Noell S, Planken SP, Rowe T, Ryan K, Smith GJI, Solowiej JE, Steppan CM, Taggart B. 2020. Discovery of Ketone-Based Covalent Inhibitors of Coronavirus 3CL Proteases for the Potential Therapeutic Treatment of COVID-19. J Med Chem 63:12725– 12747.

39. Chan HTH, Moesser MA, Walters RK, Malla TR, Twidale RM, John T, Deeks HM, Johnston-Wood T, Mikhailov V, Sessions RB, Dawson W, Salah E, Lukacik P, Strain-Damerell C, Owen CD, Nakajima T, Świderek K, Lodola A, Moliner V, Glowacki DR, Spencer J, Walsh MA, Schofield CJ, Genovese L, Shoemark DK, Mulholland AJ, Duarte F, Morris GM. 2021. Discovery of SARS-CoV-2 Mpro peptide inhibitors from modelling substrate and ligand binding. Chem Sci 12:13686–13703.

40. Fu L, Ye F, Feng Y, Yu F, Wang Q, Wu Y, Zhao C, Sun H, Huang B, Niu P, Song H, Shi Y, Li X, Tan W, Qi J, Gao GF. 2020. Both Boceprevir and GC376 efficaciously inhibit SARS-CoV-2 by targeting its main protease. 1. Nature Communications 11:4417.

41. Qiao J, Li Y-S, Zeng R, Liu F-L, Luo R-H, Huang C, Wang Y-F, Zhang J, Quan B, Shen C, Mao X, Liu X, Sun W, Yang W, Ni X, Wang K, Xu L, Duan Z-L, Zou Q-C, Zhang H-L, Qu W, Long Y-H-P, Li M-H, Yang R-C, Liu X, You J, Zhou Y, Yao R, Li W-P, Liu J-M, Chen P, Liu Y, Lin G-F, Yang X, Zou J, Li L, Hu Y, Lu G-W, Li W-M, Wei Y-Q, Zheng Y-T, Lei J, Yang S. 2021. SARS-CoV-2 Mpro inhibitors with antiviral activity in a transgenic mouse model. Science 371:1374–1378.

42. MacDonald EA, Frey G, Namchuk MN, Harrison SC, Hinshaw SM, Windsor IW. 2021. Recognition of Divergent Viral Substrates by the SARS-CoV-2 Main Protease. ACS Infect Dis 7:2591–2595.

43. Hamilton KO, Backstrom G, Yazdanian MA, Audus KL. 2001. P-glycoprotein efflux pump expression and activity in Calu-3 cells. J Pharm Sci 90:647–658.

44. Iketani S, Mohri H, Culbertson B, Hong SJ, Duan Y, Luck MI, Annavajhala MK, Guo Y, Sheng Z, Uhlemann A-C, Goff SP, Sabo Y, Yang H, Chavez A, Ho DD. 2023. Multiple pathways for SARS-CoV-2 resistance to nirmatrelvir. 7944. Nature 613:558–564.

45. Zhou Y, Gammeltoft KA, Ryberg LA, Pham LV, Tjørnelund HD, Binderup A, Duarte Hernandez CR, Fernandez-Antunez C, Offersgaard A, Fahnøe U, Peters GHJ, Ramirez S, Bukh J, Gottwein JM. 2022. Nirmatrelvir-resistant SARS-CoV-2 variants with high fitness in an infectious cell culture system. Science Advances 8:eadd7197.

