## Supplementary_information for "Identification of novel and potent inhibitors of SARS-CoV-2 main protease from DNA-encoded chemical libraries"

**Table of Contents:** **Page**

Supplementary table 1: Data collection and refinement statistics for three SARS-Cov2 -ligand complexes 2

Supplementary table 2: Metabolic stability of SLL11 and SLL12 2

Supplementary Figure 1: Avi-tagged SARS CoV-2 main protease efficiently immobilized to paramagnetic beads 3

Supplementary Figure 2: Activity of avi-tagged $M^{pro}$ coupled beads 3

Supplementary Figure 3: Off-target activity of compounds selected from DELopen 4

Supplementary figure 4: Comparison of binding modes of compounds SLL12 and MP9 4

Supplementary Figure 5: Dose response curves of compounds tested for the SAR study 5

Supplementary Figure 6: Comparison of the binding poses of the compounds SLL11 and nirmatrelvir. 6

Supplementary methods 1: Expression and purification of SARS-CoV-2 3CL protease for enzymatic assays 6-7

Supplementary methods 2: Expression and Purification Protocol of Avi-tagged $M^{pro}$ 7-10

Supplementary methods 3: Expression and purification of SARS-CoV-2 3CL protease for co-crystallization 10

Supplementary methods 4: Mpro Protease Activity Assay (Used for data shown in Table 1) 10

Supplementary methods 5: Counter Assay Screening 10

*Supplementary tables:*

Supplementary Table 1: Data collection and refinement statistics for three SARS-Cov2 $M^{Pro}$-ligand complexes

| | SLL11 | SLL12 | MP9 |
|---|---|---|---|
| **Data collection** | | | |
| Space group | P1 | P1 | P1 |
| Cell dimensions | | | |
| *a*, *b*, *c* (Å) | 69.81 85.74 93.91 | 69.83 85.56 93.67 | 69.46 85.05 93.90 |
| α, β, γ (°) | 79.25 87.44 83.86 | 78.96 87.18 83.17 | 79.50 87.62 84.14 |
| Resolution (Å) | 46.1-2.11 (2.31-2.11) | 43.9-2.25 (2.44-2.25) | 46.3-2.54 (2.76-2.53) |
| $R_{merge}$ | 0.129 (0.797) | 0.157 (0.683) | 0.187 (0.628) |
| $I / \sigma I$ | 7.3 (1.6) | 6.7 (1.8) | 4.6 (1.4) |
| Completeness spherical (%) | 63.1 (13.3) | 58.8 (13.9) | 60.0 (13.5) |
| Completeness ellipsoidal (%) | 89.1 (52.8) | 86.0 (64.3) | 88.7 (60.2) |
| Redundancy | 3.6 (3.4) | 3.5 (3.4) | 3.7 (3.7) |
| **Refinement** | | | |
| Resolution (Å) | 46.2-2.11 (2.17-2.11) | 43.9-2.25 (2.31-2.25) | 46.4-2.54 (2.60-2.54) |
| No. reflections | 72 978 (423) | 55 787 (629) | 39 749 (292) |
| $R_{work}$ / $R_{free}$ | 0.21/0.26 (0.35/0.37) | 0.25/0.29 (0.29/0.31) | 0.20/0.24 (0.31/0.34) |
| No. atoms | 14 978 | 14 761 | 14 634 |
| Protein | 14 188 | 14 131 | 14 191 |
| Ligand/ion | 282/5 | 276/7 | 270/2 |
| Water | 503 | 347 | 171 |
| *B*-factors | | | |
| Protein | 33.6 | 35.2 | 34.0 |
| Ligand/ion | 27.1 | 27.0 | 29.5 |
| Water | 26.2 | 18.9 | 15.6 |
| R.m.s. deviations | | | |
| Bond lengths (Å) | 0.006 | 0.007 | 0.006 |
| Bond angles (°) | 1.278 | 1.257 | 1.218 |
| Diffraction source | MAXIV/BioMAX | MAXIV/BioMAX | DLS i04 |
| PDB ID | 9EO6 | 9EOR | 9EOX |

Supplementary table 1: Metabolic stability of SLL11 and SLL12

| | **Human liver microsomes** | | **Mouse liver microsomes** | |
|---|---|---|---|---|
| **Compound** | **In vitro half time (min)** | **Clint (µl/min/mg)** | **In vitro half time (min)** | **Clint (µl/min/mg)** |
| SLL11 | 10 | 139 | 9 | 154 |
| SLL12 | 11 | 131 | 8 | 170 |

*Supplementary figures:*

**Supplementary Figure 1: Avi-tagged SARS CoV-2 main protease efficiently immobilized to paramagnetic beads.**

Immobilization of $M^{pro}$-Avi to paramagnetic streptavidin T1 beads was tested before DELopen selection. $M^{pro}$-Avi was efficiently captured to streptavidin T1 beads while no protein remained in the flowthrough or heated eluate samples, suggesting optimal target retention on beads during selection procedure.

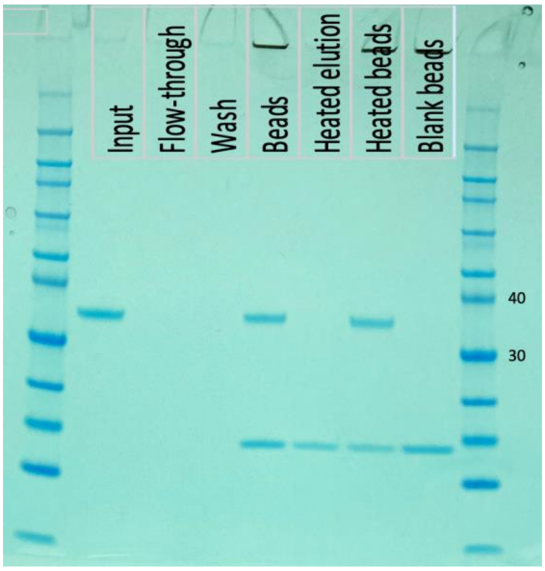

**Supplementary Figure 1:** Bead capture test of $M^{pro}$-Avi. Immobilization efficiency of C-terminally Avi-tagged $M^{pro}$ to Streptavidin T1 beads (capture to 40 ul bead slurry). Capture efficiency deduced by comparing intensity of the input and flowthrough bands around 35 kDa. Absence of protein in heated elution lane confirms that there's no release of immobilized target upon heat denaturation. Markers of 30 and 40 kDa marked.

**Supplementary Figure 2: Activity of avi-tagged $M^{pro}$ coupled beads.**

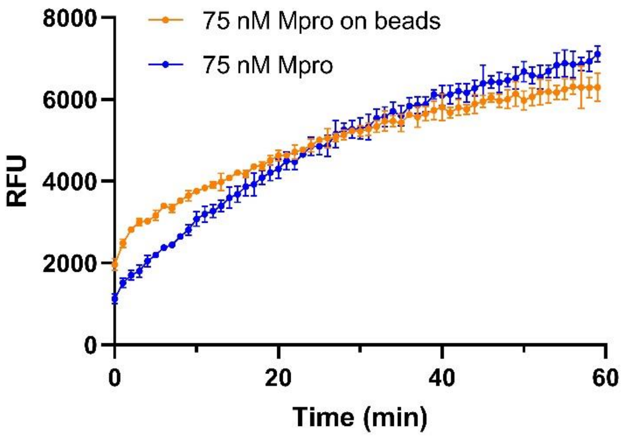

**Supplementary Figure 2:** Activity of $M^{pro}$ enzyme on beads. The $M^{pro}$ used for comparison of catalytic activity (at a final concentration of 75 nM) was obtained from the Protein Science Facility (PSF, Karolinska Institutet, Stockholm, Sweden). $M^{pro}$ activity was analyzed by detection of hydrolysis of a quenched SARS-CoV-2 $M^{pro}$ substrate (Bachem AG, Bubendorf, Switzerland). The assay was performed in 20 mM Tris, 50 mM NaCl and 0.1 mM EDTA (Merck KGaA, Darmstadt, Germany) at room temperature (pH 7.5) in Corning 3575 non-binding 384 well assay plates. $M^{pro}$, either in solution or on magnetic beads, was added to the assay plate using a 16-channel pipette (Integra ViaFlo, BergmanLabora AB, Sweden) and incubated for 15 minutes. Thereafter, the $M^{pro}$ fluorogenic substrate (stock solution in 100% DMSO) was added to the assay plate to a final concentration of 10 µM, with a Labcyte ECHO 550 non-contact dispenser and fluorescence was measured at ambient temperature in kinetic mode every minute using a PerkinElmer Envision plate reader with excitation at 340 nm and emission at 490 nm.

### Supplementary Figure 3: Off-target activity of compounds selected from DELopen.

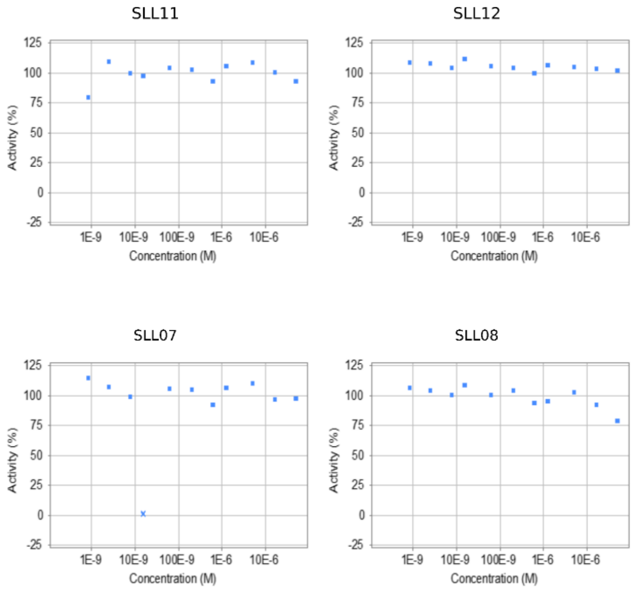

**Supplementary Figure 3**. Counter screening of compounds against human Cathepsin S enzymatic activity. Compounds SLL11, SLL12, SLL07 and SLL08 were tested at concentrations ranging from 50 to 0.00085 µM. At tested concentration no inhibitory activity was against Cathepsin S was observed.

### Supplementary figure 4: Comparison of binding modes of compounds SLL12 and MP9

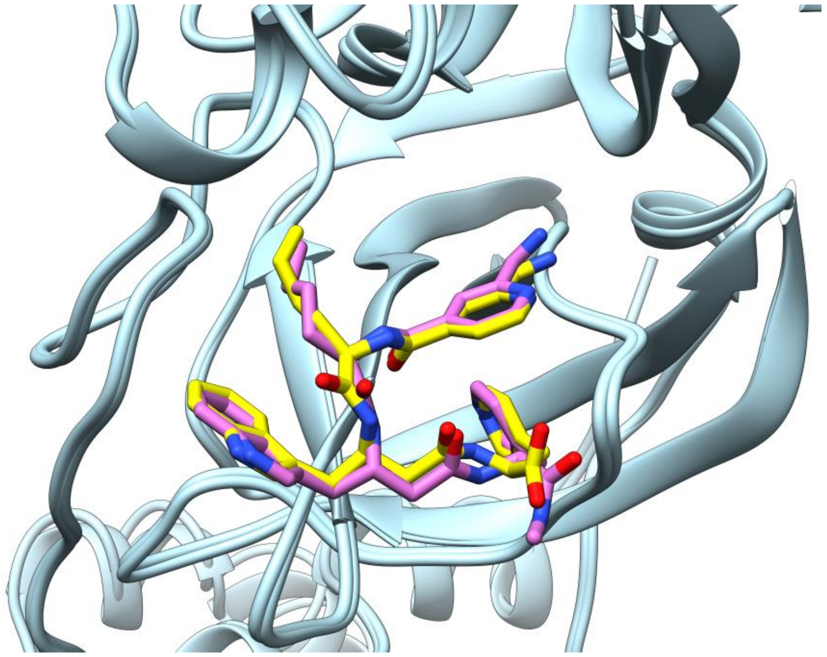

**Supplementary Figure 4.** Comparison of the binding modes of compounds SLL12 (PDB ID 9EOR) and, MP9 (PDB ID 9EOX). In compounds MP9 (shown in yellow) the C-terminal methyl amine group present in SLL12 (shown in purple) was substituted with carboxylic acid. The substitution had no effect on the compound binding mode which remains unchanged. The same dynamics likely applies also to compounds SLL11 and MP1.

### Supplementary Figure 5: Dose response curves of compounds tested for the SAR study.

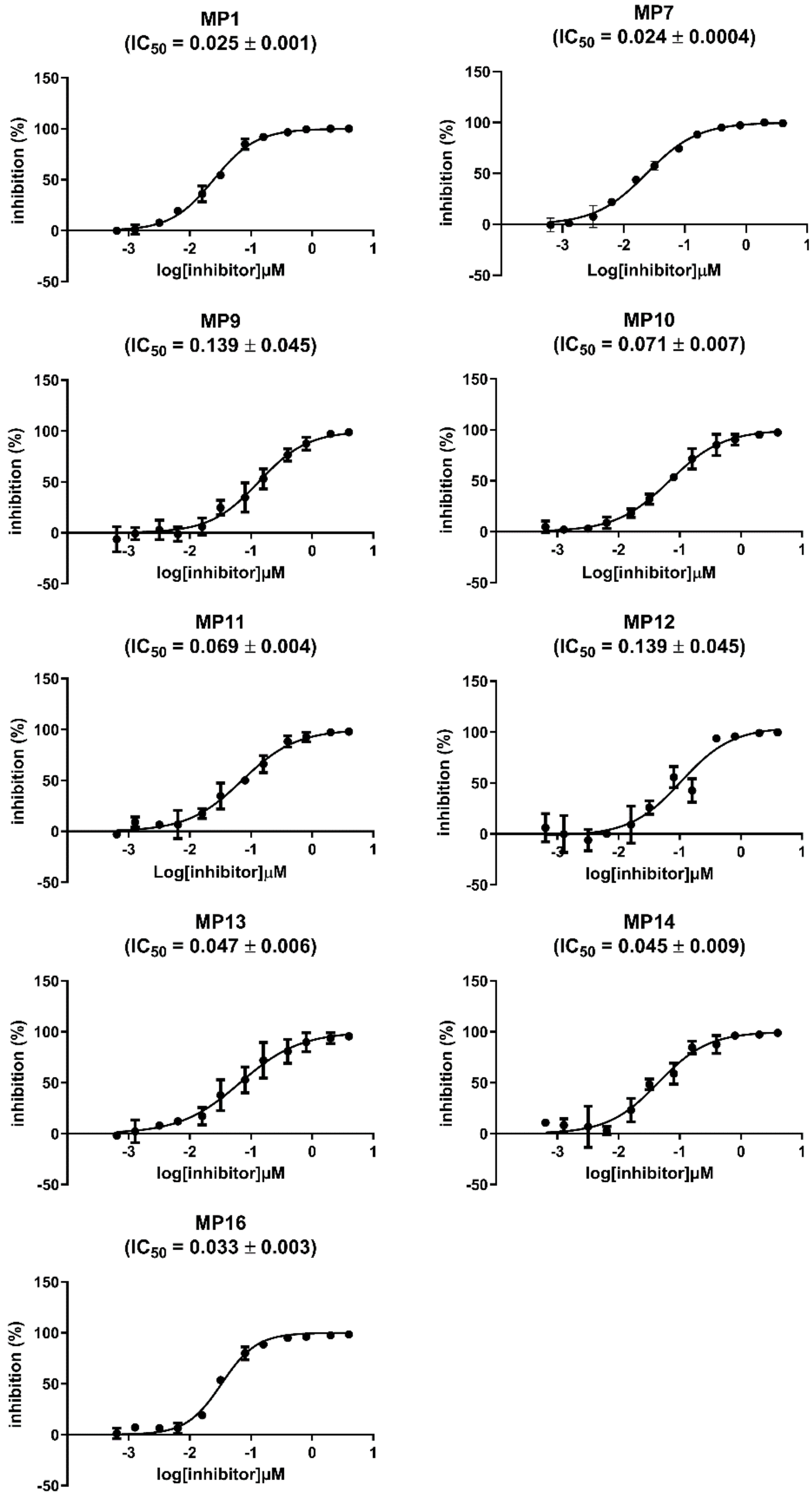

**Supplementary Figure 5.** Dose response curves and $IC_{50}$ of compound MP1 and analogues that inhibited $M^{pro}$ by 80% or more during the first screening performed at 1μM concentration. $IC50_s$ are reported as average from two independent experiments ± SEM.

**Supplementary Figure 6: Comparison of the binding poses of the compounds SLL11 and nirmatrelvir.**

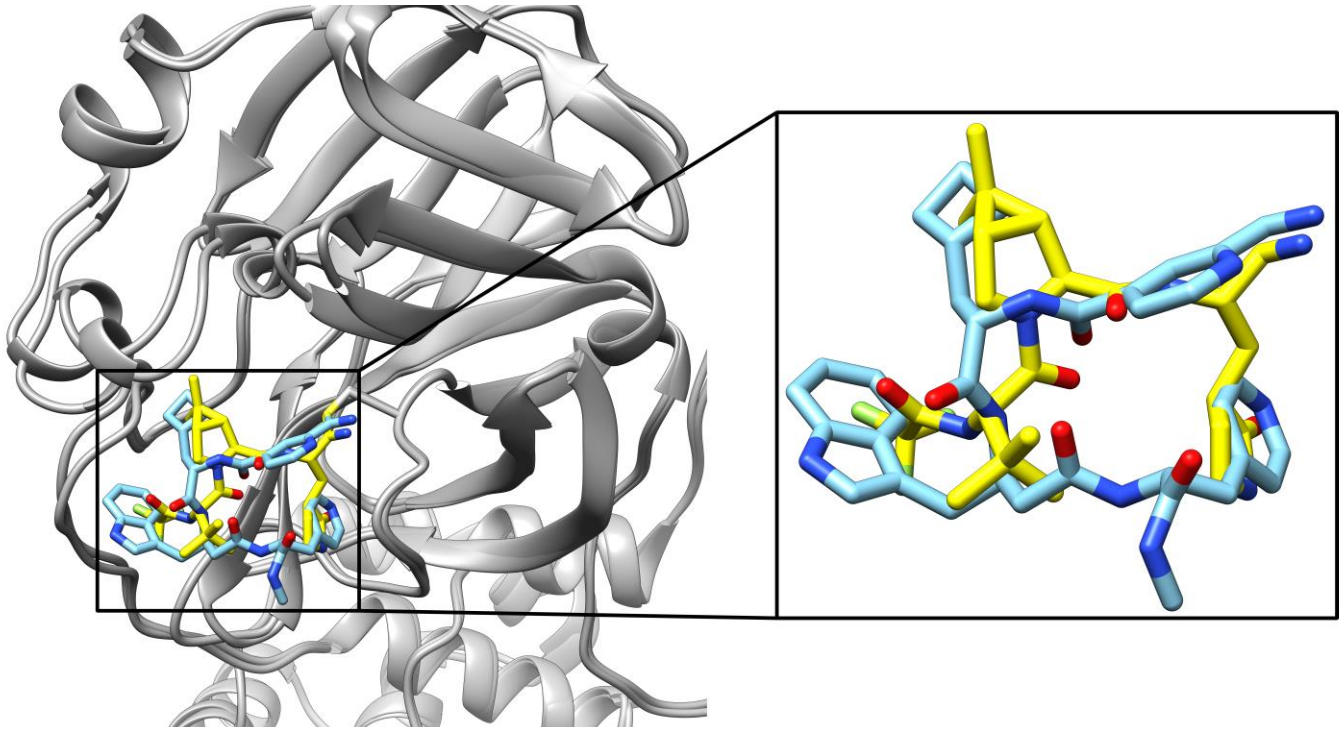

**Supplementary Figure 6.** Comparison of the binding poses of the compounds SLL11 and nirmatrelvir. Compound SLL11 binds to M$^{pro}$ with reversed orientation (C-terminus to N-terminus) assuming the conformation of a D-peptide.

*Supplementary methods:*

**1.Expression and purification of SARS-CoV-2 3CL protease for enzymatic assays**

SARS-CoV-2 3CL protease was produced by adopting a published construct used for the expression of SARS-CoV 3CL protease (Xue et al., 2007), containing nucleotide sequences corresponding to residues S1-Q306 (Chinese isolate, NCBI accession number YP_009725301). In this construct the 3CL protease is flanked by an N-GST (Glutathione S-transferases) tag and a C-HIS tag containing six histidine residues. The N-GST tag is cleaved directly by the SARS-CoV-2 protease thanks to the incorporation of the sequence of the N-terminus autocleavage site of SARS-CoV-2 3CL protease. While between the C-His tag and SARS-CoV-2 protease gene is present a cleavage site recognized by the Human rhinovirus 3C protease.

The vector (pGEX-6P-1) containing SARS-CoV-2 protease sequenced (kindly provided by Petra Lukacik, Frank von Delft and Martin Walsh) was transformed into *E. coli* BL21 (DE3)-T1R competent cells carrying the pRARE2 plasmid. A starter culture in L-Broth media (Formedium, Norfolk, UK) supplemented with carbenicillin (100 µg/ml) was inoculated from fresh transformants. The culture was grown at 37°C over the day until an $OD_{600}$ of 1.5-2 was reached. The starter culture was then used to inoculate Auto Induction Media (AIM) Terrific Broth base with trace elements (Formedium, Norfolk, UK) supplemented with 1% glycerol and carbenicillin (100 µg/ml). When the $OD_{600}$ reached 2, the cultures were shifted to 18 °C and the protein expression was continued overnight for 13 hours. Cells were harvested by centrifugation (10 min at 4500 × *g*, 4 °C), re-suspended in IMAC lysis buffer (50 mM Tris, 300 mM NaCl, pH 8.0) additionally containing Benzonase nuclease (10 µl/1.5 liter culture, 250 U/µl, E1014, Merck, Darmstadt, Germany), and disrupted by sonication (4s/4s 3 min, 80% amplitude, Sonics Vibracell-VCX750, Sonics & Materials Inc., Newtown, CT, US). Lysates were centrifuged at 49,000 × *g* for 20 min at 4 °C. The supernatants were filtered (Corning bottle-top vacuum filter, 0.45 µm, Corning, NY, USA) and imidazole was added to a final concentration of 10 mM before loading on the ÄKTA Xpress protein purification system (Cytiva, Little Chalfont, UK). Protein purification was performed using an IMAC HisTrap HP 5 ml column (Cytiva, Little

Chalfont, UK). The column was washed with wash buffer (50 mM Tris, 300 mM NaCl, 25 mM imidazole, pH 8) and the bound protein was eluted with elution buffer (50 mM Tris, 300 mM NaCl, 500 mM imidazole, pH 8). The eluted protein was then treated with HRV 3C protease (1 µg/500 µg target protein, SAE0045, Merck, Darmstadt, Germany) in order to remove the C-His tag and the buffer was at the same time exchanged by dialysis (dialysis buffer 50 mM Tris, 300 mM NaCl, 0.5 mM TCEP and 0.5 mM DTT, pH 8) with a dialysis cassette (Slide-A-Lyzer Dialysis Cassette, 10K MWCO, 3mL, Thermo Fisher Scientific, Waltham, MA, USA) over night at 4 °C. Untagged SARS-CoV-2 3CL protease was purified by reverse IMAC purification using a HisTrap 1 ml column (Cytiva, Little Chalfont, UK), the same wash and elution buffers described above were used and the flow through was collected. The reverse IMAC purification was followed by a size exclusion chromatography step using a HiLoad 16/60 Superdex 200 preparative grade column (Cytiva, Little Chalfont, UK) equilibrated with gel filtration buffer (50 mM Tris, 300 mM NaCl, pH 8) and connected to a ÄKTA Express purification system. Fractions containing the target protein were examined by SDS PAGE, pooled together, concentrated with Vivaspin® 20 ml centrifugal concentrators (10 kDa MWCO, Sartorius, Goettingen, Germany) at 4000 × *g* and the buffer was at the same time exchanged to the final storage buffer (20 mM HEPES, 50 mM NaCl, pH 7.5). The protein was finally flash frozen in liquid nitrogen and stored at −80 °C.

### 2. Expression and Purification Protocol of Avi-tagged M$^{pro}$ (for affinity selection of "DELopen" DNA encoded chemical library)

#### 2.1-Construct details

M$^{pro}$ was amplified with primers that would incorporate a C-terminal Avi-Tag, in place of the existing 6His tag. This was introduced into BamHI/XhoI cut pGEX-6P-1 by In-Fusion cloning. The resultant construct is GST-3C-✂M$^{pro}$-3C-AviTag, with the GST-3C portion being autocleaved by M$^{pro}$ during expression (✂) and the terminal 3C site incorportaing the native C-terminal residues of M$^{pro}$. Purification is achieved by *in vivo* biotinylation during expression and a Streptavidin Mutein Matrix (Roche) pull down.

#### 2.2-DNA sequence:

ATGTCCCCTATACTAGGTTATTGGAAAATTAAGGGCCTTGTGCAACCCACTCGACTTCTTTTGGAATATCTTGAA
GAAAAATATGAAGAGCATTTGTATGAGCGCGATGAAGGTGATAAATGGCGAAACAAAAAGTTTGAATTGGGTTT
GGAGTTTCCCAATCTTCCTTATTATATTGATGGTGATGTTAAATTAACACAGTCTATGGCCATCATACGTTATATAG
CTGACAAGCACAACATGTTGGGTGGTTGTCCAAAAGAGCGTGCAGAGATTTCAATGCTTGAAGGAGCGGTTT
TGGATATTAGATACGGTGTTTCGAGAATTGCATATAGTAAAGACTTTGAAACTCTCAAAGTTGATTTTCTTAGCAA
GCTACCTGAAATGCTGAAAATGTTCGAAGATCGTTTATGTCATAAAACATATTTAAATGGTGATCATGTAACCCAT
CCTGACTTCATGTTGTATGACGCTCTTGATGTTGTTTTATACATGGACCCAATGTGCCTGGATGCGTTCCCAAA
ATTAGTTTGTTTTAAAAAACGTATTGAAGCTATCCCACAAATTGATAAGTACTTGAAATCCAGCAAGTATATAGCA
TGGCCTTTGCAGGGCTGGCAAGCCACGTTTGGTGGTGGCGACCATCCTCCAAAATCGGATCTGGAAGTTCTG
TTCCAGGGGCCCCTGGGATCCGCTGTCTTACAGTCCGGTTTCCGTAAGATGGCCTTCCCGTCAGGCAAAGTG
GAAGGCTGTATGGTGCAAGTAACGTGCGGGACCACAACGTTGAACGGTTTGTGGTTAGATGATGTGGTTTACT
GTCCACGTCATGTTATCTGTACAAGTGAGGACATGCTTAATCCAAATTATGAGGACTTGTTGATCCGCAAAAGC
AACCATAATTTCCTGGTACAGGCAGGCAACGTTCAGTTACGTGTCATTGGTCATTCAATGCAAAACTGCGTGTT
GAAGCTGAAAGTAGATACAGCCAACCCGAAAACCCCGAAATATAAATTTGTCCGTATTCAGCCAGGCCAGACC
TTTTCGGTTCTGGCGTGCTACAACGGTAGCCCATCTGGGGTCTACCAGTGCGCTATGCGTCCTAACTTTACAA
TTAAGGGCAGTTTCTTGAACGGTAGCTGCGGAAGCGTTGGCTTTAATATTGACTACGATTGTGTGTCATTTTGC
TATATGCACCACATGGAGTTACCCACTGGAGTTCACGCGGGAACTGACCTGGAGGGGAACTTCTATGGGCCA
TTTGTGGATCGCCAGACGGCGCAGGCGGCCGGAACGGATACTACTATTACCGTGAATGTCCTTGCTTGGTTAT
ACGCGGCCGTCATTAACGGTGACCGTTGGTTTTTAAACCGTTTCACCACGACCCTTAATGATTTTAATTTGGTG
GCTATGAAGTATAACTACGAACCCCTGACGCAGGACCACGTAGATATTTTGGGGCCGCTGTCGGCACAGACG
GGAATTGCAGTGTTAGATATGTGTGCTTCATTGAAAGAGTTGTTACAGAACGGTATGAATGGACGCACAATTTT
GGGATCAGCATTATTAGAGGATGAGTTTACTCCGTTTGATGTTGTGCGTCAGTGCTCGGGTGTAACTTTCCAG
GGGCCGGGCCTGAATGATATTTTCGAGGCACAGAAAATTGAATGGCATGAGTGA

**2.3-Protein sequence:**

MSPILGYWKIKGLVQPTRLLLEYLEEKYEEHLYERDEGDKWRNKKFELGLEFPNLPYYIDGDVKLTQSMAIIRYIADK
HNMLGGCPKERAEISMLEGAVLDIRYGVSRIAYSKDFETLKVDFLSKLPEMLKMFEDRLCHKTYLNGDHVTHPDFM
LYDALDVVLYMDPMCLDAFPKLVCFKKRIEAIPQIDKYLKSSKYIAWPLQGWQATFGGGDHPPKSDLEVLFQGPLG
SAVLQ✂SGFRKMAFPSGKVEGCMVQVTCGTTTLNGLWLDDVVYCPRHVICTSEDMLNPNYEDLLIRKSNHNFLV
QAGNVQLRVIGHSMQNCVLKLKVDTANPKTPKYKFVRIQPGQTFSVLACYNGSPSGVYQCAMRPNFTIKGSFLNG
SCGSVGFNIDYDCVSFCYMHHMELPTGVHAGTDLEGNFYGPFVDRQTAQAAGTDTTITVNVLAWLYAAVINGDR
WFLNRFTTTLNDFNLVAMKYNYEPLTQDHVDILGPLSAQTGIAVLDMCASLKELLQNGMNGRTILGSALLEDEFTPF
DVVRQCSGVTFQGPGLNDIFEAQKIEWHE

GST, 3C sites, $M^{pro}$, $M^{pro}$ autocleavage site (✂), Avi-Tag

**2.4- Expression method:**

Expression Strain:

Rosetta-BirA: BL21(DE3)-pRARE-2-pCDF-BirA (Chl/Strep)

AIM TB medium:

55.85 g Formedium AIM TB, including trace elements

10 mL glycerol

Topped up to 1 L with MilliQ water

Autoclaved to sterilise.

10 mM Biotin:

For each litre of culture 48.8 mg D-Biotin (IBA-LifeSciences) were weighed out and made up to 20 mL with 10 mM Bicine, pH 8.3, buffer (pre-heated for 30 seconds at full power in the microwave). This was vortexed to fully dissolve the Biotin and then filtered through a 0.22 µm syringe filter to sterilise.

Growth conditions:

A streak of colonies from each Rosetta-BirA transformation plate was inoculated into 1 mL LB medium, containing 100 µg/mL carbencillin and 50 µg/mL Streptomycin, and grown at 37°C 650 rpm for 1 hour. This was then inoculated into 10 mL LB, containing 100 µg/mL carbencillin and 50 µg/mL Streptomycin, and grown at 37°C 200 rpm for a further 2 hours. This was finally inoculated into 100 mL LB medium, containing 100 µg/mL carbencillin and 50 µg/mL Streptomycin, and grown at 37°C 200 rpm for 5 hours. Finally 10 mL of the starter culture were inoculated into each 2.5 L baffled flask of pre-warmed 1 L AIM TB, containing 100 µg/mL carbenicillin, 50 µg/mL Streptomycin and 20 mL of 10 mM Biotin solution. This was grown at 37°C 200 rpm for 5 hours at which point the temperature was reduced to 18°C overnight. The cells were harvested by centrifugation at 4,500 rpm for 30 minutes at 4°C.

**2.5-Purification:**

Buffers:

| | |
|---|---|
| Lysis Buffer: | 50 mM Tris, pH 8.0 |
| | 300 mM NaCl |

Elution Buffer 1: 50 mM Tris, pH 8.0

300 mM NaCl

10 mM Biotin

Elution Buffer 2: 50 mM Tris, pH 8.0

300 mM NaCl

50 mM Biotin

(+26 mM NaOH (130 µL 10 mM NaOH per 50 mL) to get Biotin into solution)

Mutein Wash Buffer: 100 mM potassium phosphate buffer, pH 7.2

150 mM NaCl

Mutein Equilibration Buffer: 100 mM potassium phosphate buffer, pH 7.2

150 mM NaCl

400 mM Ammonium sulfate

50% Streptavidin Mutein Matrix: Streptavidin Mutein Matrix (Roche)

Washed twice with Mutein Wash Buffer

Washed three times in Mutein Equilibration Buffer

Resuspended to 50% in Mutein Equilibration Buffer

Purification protocol:

A 1L pellet (~25 g cells) was thawed at room temperature and decanted into a beaker.

Lysis Buffer was added at a ratio of 30 mL per 10 g of cells and supplemented with 3 µg Benzonase (do not add protease inhibitors). This was stirred at 4 °C until the cell pellet had fully resuspended.

The cells were lysed by sonication for 5 minutes on ice, with 15 seconds on and 15 seconds off (a cell disruptor may also be used).

The lysate was decanted into 50 mL JA 25.50 centrifuge tubes and centrifuged for 1 hour at 50,000 xg.

The cleared lysate was carefully decanted off into 50 mL falcon tubes and a total of 5 mL (bed volume) of the Streptavidin Mutein Matrix split across the tubes. These were then placed on a rotary mixer for 1 hour at 4°C.

The mixture was transferred into a BioRad Econo-Column, the flow-through discarded.

The resin was washed 3 times with 40 mL Lysis Buffer.

5 mL of Elution Buffer 1 were applied and incubated for 10 minutes before eluting.

The remainder of the protein was then eluted with another 5 mL of Elution Buffer 1 and two additions of Elution Buffer 2.

The elutions were analysed by SDS-PAGE (example shown below) and the appropriate fractions pooled.

The sample was concentrated down to 5 mL using an Amicon Ultra-15 10 kDa MWCO concentrator, passed through a 0.22 um filter onto an S200 16/60 HiLoad Gel Filtration column run in Lysis Buffer. This step removes the Biotin and contaminating GST only band.

The GFC fractions were analysed by SDS-PAGE, appropriate fractions pooled ($M^{pro}$ runs as a dimer) and concentrated to the desired final concentration before snap freezing in liquid nitrogen.

**3. Expression and purification of SARS-CoV-2 3CL protease for co-crystallization experiments**

SARS-CoV-2 3CL protease for co-crystallization experiments was produced using the same protocol as for the protein used for enzymatic assays except for the following steps. After the first IMAC purification step the protein was further purified by size exclusion chromatography (SEC) using a HiLoad 16/60 Superdex 200 preparative grade column (Cytiva, Little Chalfont, UK) pre-equilibrated with gel filtration buffer A (50 mM Tris, 300 mM NaCl, pH 8.0). Selected fractions were analyzed by SDS-PAGE and the protein containing fractions were treated with HRV 3C protease (own production, 1:500 molar ratio) overnight at 4°C in gel filtration buffer A supplemented with 0.5 mM DTT. The cleaved SARS-CoV-2 3CL protease sample was supplemented with 20 mM imidazole and purified by reverse IMAC purification using an IMAC HisTrap HP 1 ml column (Cytiva, Little Chalfont, UK) equilibrated with wash buffer (50 mM Tris, 300 mM NaCl, 25 mM imidazole, pH 8). The flow through was collected and the protein was further purified by a second SEC step using the same column as above pre-equilibrated with gel filtration buffer B (20 mM HEPES, 50 mM NaCl, pH 7.5). Fractions containing the target protein were examined by SDS PAGE, combined, and concentrated with Vivaspin® 20 ml centrifugal concentrators (10 kDa MWCO, Sartorius, Goettingen, Germany) at 4000 × g, 4 °C. The protein was finally flash frozen in liquid nitrogen and stored at −80 °C.

**4. Mpro Protease Activity Assay (Used for data shown in Table 1).**

An internally quenched fluorogenic substrate for SARS-CoV-2 Mpro (DABCYL-Lys-HCoV-SARS Replicase Polyprotein 1ab (3235−3246)-Glu-EDANS trifluoroacetate salt, >95% pure) was custom synthesized and obtained from Bachem AG, Switzerland. The Mpro protein used for catalytic activity assays was obtained from the Protein Science Facility (PSF, Karolinska Institutet, Stockholm, Sweden) and is described in a prior section. All test compounds were dissolved to 10 mM stocks in 100% DMSO (Merck KGaA, Darmstadt, Germany) and transferred to Echo LDV source plates (Labcyte, Inc. CA). Mpro activity was analyzed by detection of hydrolysis of an internally quenched SARS-CoV-2 Mpro substrate. The assay was performed in 20 mM Tris, 50 mM NaCl, 0.01% Triton X-100 and 0.1 mM EDTA (Merck KGaA, Darmstadt, Germany) at room temperature (pH 7.5). Compounds were transferred in an 11-point concentration series (1:3 dilutions, starting concentration 50 µM) with an Echo 550 non-contact dispenser (Labcyte, Inc., USA) to Corning 3575 non-binding 384 well assay plates. Mpro (75 nM final concentration) was added to the assay plate using a 16-channel pipet (Integra ViaFlo, Bergman Labora AB, Sweden) and shaken for 15 min at 1000 rpm in an Eppendorf Mixmate. After a pulse centrifugation, the Mpro fluorogenic substrate (stock solution at 5 mM in 100% DMSO) was added to the assay plate to a final concentration of 10 µM, thus contributing with 0.2% DMSO in final assay, with a Labcyte Echo 550 noncontact dispenser. After 10 min incubation with shaking at 1000 rpm in an Eppendorf Mixmate and a pulse centrifugation, fluorescence was measured in a PerkinElmer Envision plate reader at ambient temperature with excitation at 340 nm and emission at 490 nm.

**5. Counter Assay Screening.**

The effect of selected compounds on the activity on human cathepsin S was determined with the SensoLyte 440 cathepsin S Assay Kit (Anaspec, Inc.) used according to the manufacturer's recommendations.
